# *Foxn3* is part of a transcriptional network that regulates primary cilia in the developing retina

**DOI:** 10.1101/2024.10.28.620684

**Authors:** Huanqing Zhang, Thejas Nair, Fan Meng, David L. Turner

## Abstract

Analysis of retinas from mice with a targeted disruption of the gene encoding the forkhead transcription factor Foxn3 revealed additional displaced amacrine interneurons and retinal astrocytes in the inner plexiform and ganglion cell layers, as well as ectopic primary cilia on bipolar and amacrine interneurons. Foxn3 is a transcriptional repressor and numerous genes linked to cilia structure or assembly were upregulated in embryonic retinas without *Foxn3*, including Foxj1, a forkhead transcription factor required for motile cilia. CUTCRUN analysis revealed that many of the upregulated retinal genes were bound by the Foxn3 and Rfx3 proteins. A short hydrophobic motif (LXXLXWL) shared by the Foxj1, Foxn4, and Foxn3 proteins was found to be required for association with the Rfx3 protein and for transcriptional repression by Foxn3, as well as for full transcriptional activation by Foxn4 with Rfx3. AlphaFold 3 predicts an interaction between the shared hydrophobic motif and the Rfx3 dimerization domain. Mutations in Rfx3 at the predicted interaction site disrupted association of Rfx3 with Foxn3 or Foxn4. These results reveal a new layer of transcriptional regulation of genes required for cilia, with Foxn3 functioning as a repressor of cilia genes and limiting primary cilia formation in the developing retina.

## Introduction

Cilia are thin, filamentous, membrane-enclosed structures with a microtubule core, present on cells in many eukaryotes (reviewed in Derderian et al., 2023). Motile cilia can direct fluid flow or drive cellular movement, while primary cilia (also known as sensory cilia) mediate reception of various extracellular signals, including Sonic Hedgehog (SHH), TGFβ, PDGF-A, Wnt, and Notch. Photoreceptor outer segments are specialized primary cilia used for phototransduction (Chen et al., 2021). In the mature retina, retinal ganglion cells, amacrine interneurons, and Müller glia have been reported to have primary cilia in the mouse, while most bipolar cells do not (Kowal et al., 2022; Ning et al., 2021). During development, retinal progenitor cells have primary cilia located at the outer surface of the retina (Burnett et al., 2017; Dubaic et al., 2023). The assembly and function of cilia involve proteins produced by hundreds of genes; ciliopathies can arise from mutations in many of these genes. Ciliopathies frequently affect photoreceptors, lead to retinal degeneration, and impact vision (Chen et al., 2021). In addition, defects in primary cilia or in the signaling pathways located in primary cilia can alter cell fates or retinal structure during development (Cwinn et al., 2011; Dilan et al., 2019; Fiore et al., 2020; Wang et al., 2005; Zhang and Yang, 2001).

Members of the forkhead and regulatory factor X (RFX) transcription factor protein families regulate genes associated with cilia in multiple organisms. Forkhead proteins bind DNA via the forkhead domain and have multiple roles in development (reviewed in Herman et al., 2021). In vertebrates, the forkhead protein Foxj1 regulates genes encoding components of motile cilia (Choksi et al., 2014; Patir et al., 2020; Vij et al., 2012; Yu et al., 2008) and cooperates with the Rfx2-3 transcription factors (Didon et al., 2013; Lemeille et al., 2020). Foxn4 functions with Foxj1 to allow the generation of multiciliated cells and both are expressed in ciliated ependymal cells of the central nervous system (CNS (Campbell et al., 2016; Quigley and Kintner, 2017; Zheng et al., 2025). A *Foxj1* homolog is necessary for neurogenesis during zebrafish retinal regeneration after injury, but it is not required for retinal development in zebrafish (Lyu et al., 2023).

Over 40 genes encode forkhead proteins in mice or humans; mutations in forkhead genes are associated with genetic diseases in both organisms (Herman et al., 2021). The FOXN subfamily of forkhead proteins has four members in mice and humans. *Foxn1* is required for T-cell development (Boehm et al., 2003; Žuklys et al., 2016). *Foxn4* is required for the formation of most or all horizontal and many amacrine interneurons in the retina (Li et al., 2004) and plays an essential role in developmental timing in the retina, promoting interneuron and photoreceptor formation while suppressing retinal ganglion cells (Liu et al., 2020). *Foxn4* also functions in spinal cord interneuron development (Misra et al., 2014), and has been linked to heart development in zebrafish (Chi et al., 2008). A partial clone of human *Foxn3* was originally identified as a DNA damage checkpoint suppressor in a functional screen in yeast, while the full length Foxn3 protein functions as a transcriptional repressor in mammalian cells (Huot et al., 2014; Pati et al., 1997; Scott and Plon, 2005). *Foxn3* has been implicated in the regulation of glucose metabolism in liver (Karanth et al., 2019) and the regulation of inflammation in lungs (Yu et al., 2025; Zhu et al., 2023). Foxn2 has extensive sequence similarity to Foxn3, and both proteins have been reported to function as tumor suppressors (reviewed in Song et al., 2023). Most forkhead factors recognize one of two unrelated DNA sequences, depending on their subfamily, but Foxn3 recognizes both sequences, potentially allowing it to regulate a larger set of target genes than other forkhead factors (Nakagawa et al., 2013; Rogers et al., 2019).

We identified the *Foxn3* mRNA as a target of microRNA miR-216b in the mouse retina (Zhang et al., 2022). *Foxn3* is expressed in retinal progenitor cells as well as differentiated bipolar and cholinergic amacrine cells (Boudreau-Pinsonneault et al., 2023; Zhang et al., 2022). When overexpressed in the neonatal retina, miR-216b promotes the generation of additional amacrine cells and the loss of bipolar cells. Co-expression of Foxn3 partially reversed this effect, while inhibition of *Foxn3* generated similar phenotypes (Zhang et al., 2022). To further investigate the role of *Foxn3* in the developing retina, we analyzed retinas from *Foxn3* knockout (KO) mice (Birling et al., 2021). We observed additional amacrine cells and astrocytes in the inner plexiform layer (IPL) and ganglion cell layer (GCL) of P21 retinas from *Foxn3* KO mice, as well as ectopic primary cilia formation on bipolar and amacrine cells. In developing retinas without functional Foxn3, numerous genes involved in cilia formation and function were upregulated. We mapped genomic regions bound by the Foxn3, Foxn4, and Rfx3 proteins and found that Foxn3 binds to many genes required for cilia. We show that Foxn3 or Foxn4 can form a complex with Rfx3, and we identified specific amino acids in these proteins that are required for interaction, as well as for cooperative transcriptional activation or repression by Rfx3 with Foxn4 or Foxn3 respectively. Our results indicate that Foxn3 is an important negative regulator of cilia genes and acts to limit primary cilia formation in the retina.

## Results

### Additional and ectopic amacrine cells and astrocytes in Foxn3 KO retinas at P21

Mice heterozygous for a targeted deletion of *Foxn3* exon 3 (Birling et al., 2021) were backcrossed to generate homozygous *Foxn3* KO mice. *Foxn3* KO embryos were present at the expected frequency (25.1%; N= 183) at the end of gestation on embryonic day 19 (E19) and embryos appeared grossly normal, as previously reported (Birling et al., 2021). Nearly all *Foxn3* KO offspring died at or shortly after birth, consistent with prior reports (Birling et al., 2021; Zhu et al., 2023), but we recovered three viable *Foxn3* KO mice among mice genotyped after P7 (N=221; estimated *Foxn3* KO survival rate ∼3%). *Foxn3* KO offspring were smaller than wild-type (WT) offspring (mean±s.d.: *Foxn3* KO 4.57±1.14 gm, N=3; WT 13.96±0.47 gm, N=4; p<0.0001, unpaired t test) but appeared otherwise normal at P21. Retina sections from P21 *Foxn3* KO mice revealed that while the number of AP2α (Tfap2a)- positive amacrine cells in the inner nuclear layer (INL) did not change, displaced amacrine cells located in the GCL increased compared to WT, and ectopic amacrine cells were found within the IPL (Fig 1A-C). The number of Vsx2-positive bipolar cells and Sox9-positive Müller glia in the INL did not change (Fig 1C). However, Sox9-positive glial cells located within the IPL and GCL of the *Foxn3* KO retinas increased (Fig 1D). Sox9/Glial fibrillary acidic protein (GFAP)-positive astrocytes were present in the IPL and GCL of the *Foxn3* KO retinas, while Sox9/glutamine synthetase (Glul)-positive Muller cell bodies remained in the INL (Fig. 1E, F; Supplemental Fig. S1A, B), indicating that Sox9-positive cells in the IPL and GCL are astrocytes. GFAP-labeled processes from the astrocytes extended throughout the IPL in the Foxn3 KO retinas, in contrast to WT (Fig. 1E, F; Supplemental Fig. S1A, B). Retina sections from E19 *Foxn3* KO embryos did not show obvious changes in AP2α-positive amacrine cells or overall retinal structure (Supplemental Fig. S1C-F), indicating changes in amacrine cells at P21 likely arise during postnatal development.

**Figure 1.**
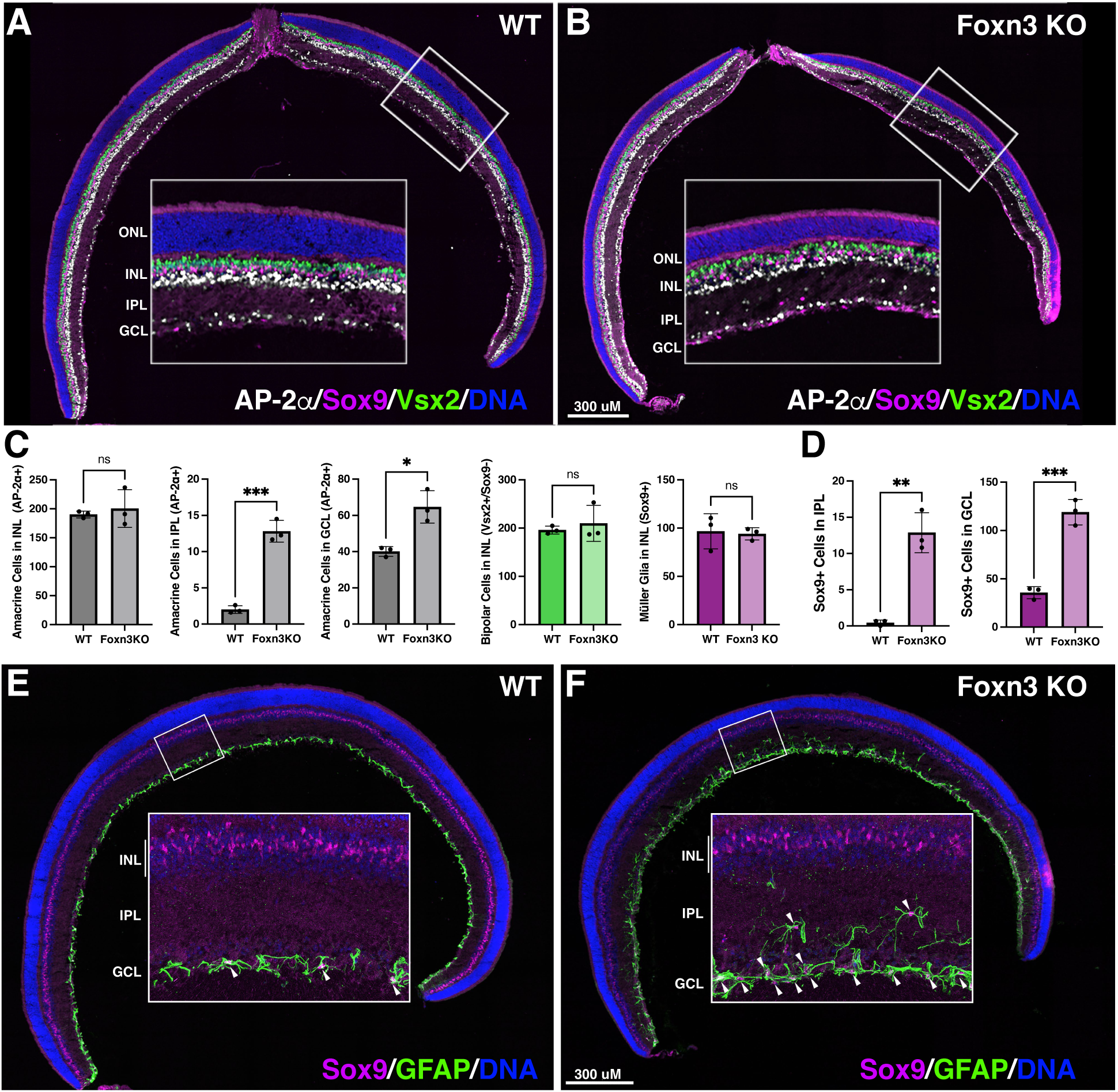
Increased and ectopic amacrine cells and astrocytes are present in Foxn3 KO retinas. Sections from WT and *Foxn3* KO P21 retinas (A,B) were analyzed by immunofluorescence; enlarged inset regions indicated. *Foxn3* KO retinas have ectopic amacrine cells (labeled with AP2α, white nuclei) and glial cells (labeled with Sox9, magenta nuclei) in the IPL (inner plexiform layer), as well as increased displaced amacrine cells and Sox9+ cells in the GCL (ganglion cell layer). In the INL (inner nuclear layer), amacrine cells, bipolar cells (labeled with Vsx2, green) and Müller glia (Sox+ nuclei in INL) were not changed. (C, D) Quantitation with unpaired t-test: *** p< 0.001, ** p< 0.01, * p< 0.05, ns = not significant. N=3 for *Foxn3* KO and WT. Counts in C are from 200um wide windows; counts in D are from complete sections (see methods). (E, F) Cells labeled by Sox9 in the IPL or GCL are retinal astrocytes that express GFAP (glial fibrillary acidic protein, green). ONL: outer nuclear layer. Nuclear DNA: blue; scale bars as indicated.

### Genes associated with cilia are upregulated in embryonic Foxn3 KO retinas

To investigate gene expression changes in the retina after loss of *Foxn3*, we used RNA-seq to compare gene expression in retinas isolated from WT and *Foxn3* KO embryos at E19 and found that 1655 genes were differentially expressed (FDR<0.01, Supplemental Table 1). Most differentially expressed genes were upregulated in the *Foxn3* KO retinas, including nearly all mRNAs that changed more than two-fold (Fig. 2A), consistent with Foxn3 repressing transcription. The mRNA for Foxj1, a transcriptional activator that regulates the gene expression program of motile cilia was upregulated ∼20-fold and several genes associated with cilia were upregulated over 100-fold (e.g., *Cfap65*, *Cfap44*, and *Cfap52*). Foxj1 protein was increased in the nuclei of the neuroblast layer in the Foxn3 KO embryonic retinas (Supplemental Fig. S2). mRNAs for Rfx1, Rfx2, and Rfx3, transcription factors linked to cilia gene expression, were significantly upregulated although each increased less than two-fold (Supplemental Table 1). The *Foxn3* gene without exon 3 expressed an mRNA with exon 2 spliced to exon 4 that was significantly upregulated in *Foxn3* KO retinas, although also with a less than two-fold increase. Strikingly, most of the top enriched GO categories for the differentially expressed genes were components of cilia and/or were associated with cilia assembly (Fig. 2B, C). 305 significantly upregulated genes are part of the ciliome, genes that contribute to cilia formation or function (Elliott et al., 2023)(Supplemental Table 1), and the fraction of ciliome genes increased among highly upregulated genes (Fig. 2D). Several genes previously reported to be repressed by Foxn3 overexpression in human cells (Huot et al., 2014) were upregulated in Foxn3 KO retinas (e.g., *Pim2*, *Dryk3*, *Iqck*, and *Ift88*).

**Figure 2.**
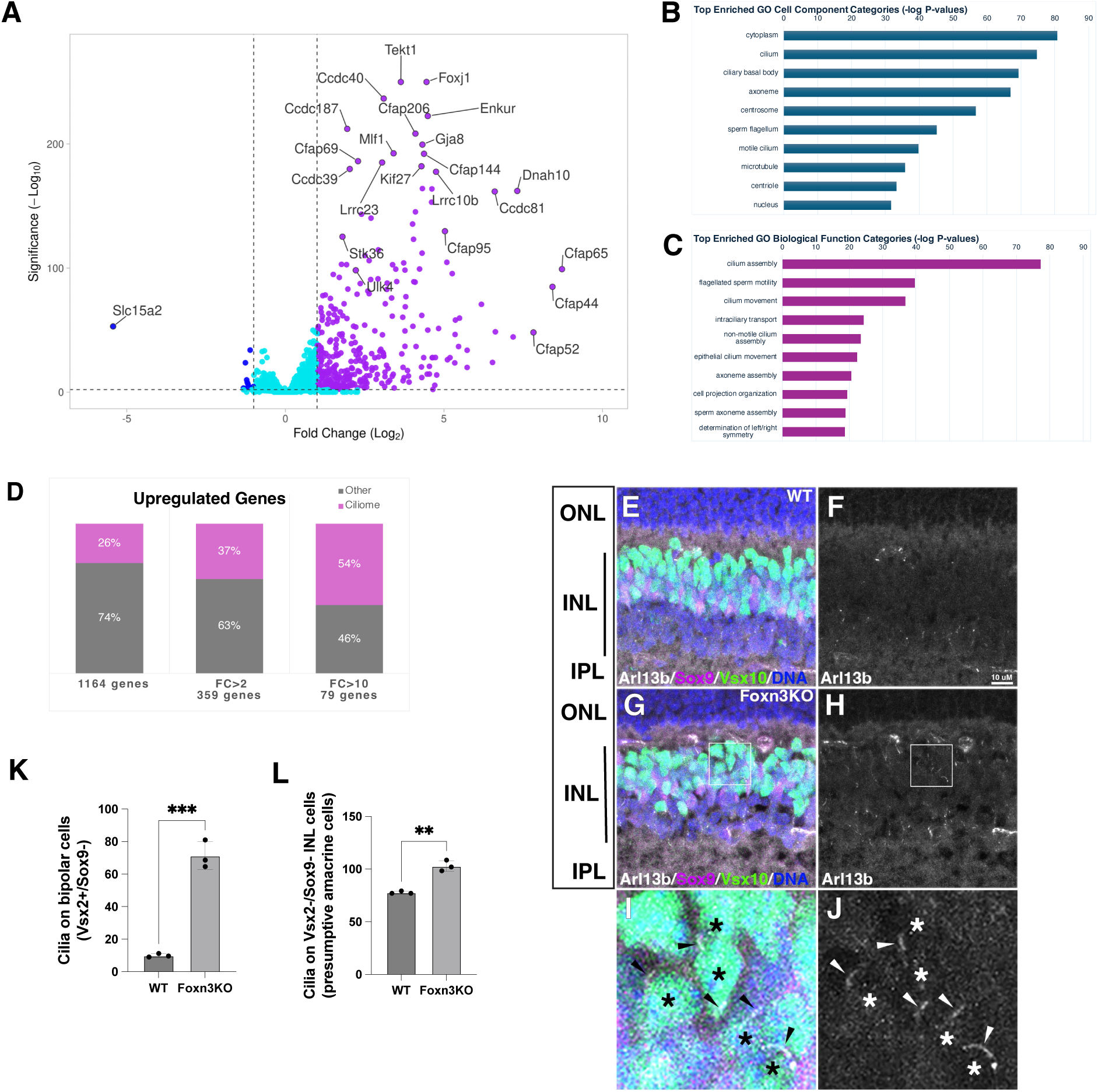
Repression of cilia genes and primary cilia in the retina by Foxn3. (A) Volcano plot showing RNA-seq analysis of E19 *Foxn3* KO retinas compared to WT retinas shows that most of 1655 differentially expressed genes (DEG) (FDR q<0.01) are upregulated in the *Foxn3* KO retinas. mRNAs significantly upregulated at least two-fold are indicated in magenta, while at least two-fold significantly downregulated mRNAs are labeled in dark blue. Labeled mRNAs in (A) are part of the ciliome, except Gja8, Lrrc10b, and Slc15a2. For Tekt1 and Foxj1 q=0; -log_10_ q was set to 250 for these two genes. (B, C) GO analysis reveals DEG are enriched for multiple GO categories related to cilia. (D) The ciliome-associated percentage of upregulated DEG increases with larger fold-change (FC). (E-K) Ectopic cilia (Arl13b, white) in the INL of *Foxn3* KO retinas. In WT retinas (E, F) few cilia are associated with bipolar cells (Vsx2-positive, green; Sox9-negative, magenta), but cilia are frequent on bipolar cells in the *Foxn3* KO retinas (G, H, K). (I, J) Enlargement of boxed regions in (G, H) showing Vsx2-positive bipolar cells (asterisks) with Arl13b-positive cilia (arrowheads) in a Foxn3 KO retina. Primary cilia on presumptive amacrine cells in the inner INL (Vsx2-negative, Sox9-negative) also increased in the Foxn3 KO (E-H, L). Nuclear DNA: blue. (I-K) N=3 for Foxn3 KO and WT; labels as in Fig.1 legend.

### Ectopic cilia in the INL of Foxn3 KO retinas

Foxn3 is expressed in retinal bipolar cells (Zhang et al., 2022), which generally lack primary cilia (Ning et al., 2021). Immunofluorescence analysis of Arl13b, a protein present in primary cilia, revealed numerous cilia on present on Vsx2 (Chx10)-expressing bipolar cells in Foxn3 KO retinas at P21, with a 7-fold increase over WT retinas (Fig. 2E-K). Arl13b labeled cilia on amacrine cells in the inner INL (Ning et al., 2021) in both genotypes, but significantly more cilia were detected in the amacrine cell layer of Foxn3 KO retinas (Fig. 2L). Many retinal progenitor cells have primary cilia located at the outer surface of the retina during retinal development (Burnett et al., 2017; Ning et al., 2021). We examined primary cilia in the retinas from *Foxn3* KO and WT mice at E19. Immunofluorescence analysis revealed variation in the expression of Ift88, a component of the intraflagellar transport (IFT) complex, among Arl13b labeled primary cilia in the Foxn3 KO retinas (Supplemental Fig. S3). The Ift88 mRNA and several other mRNAs for components of the IFT complex were upregulated two-fold or more (Supplemental Table 1) in Foxn3 KO retinas.

### Genomic binding sites for endogenous Foxn3, Rfx3, and Foxn4 in the developing retina

To determine whether genes differentially expressed in *Foxn3* KO retinas were likely to be direct targets of Foxn3, we performed CUTCRUN analysis (Skene and Henikoff, 2017) to identify genomic regions bound by the endogenous Foxn3 protein in WT mouse retinas at E16. 2794 regions with Foxn3 binding were identified (Supplemental Table 2), based on peaks of read enrichment that were consistent between two separate experiments, each with two anti-Foxn3 samples and two IgG controls. Heat maps for individual CUTCRUN libraries showed consistent signals at the identified Foxn3 binding regions (Fig. 3A). 90% of the genomic regions bound by Foxn3 overlap with mouse candidate cis-Regulatory Elements (cCREs) annotated by ENCODE (Abascal et al., 2020), with binding at promoters being most frequent (Fig. 3B). We annotated Foxn3-bound regions based on genes with the nearest transcription start site (TSS). For *Foxn3* KO retinas, 49% of the upregulated genes and 68% of the upregulated ciliome genes were linked to at least one genomic region bound by Foxn3 in the CUTCRUN analysis, indicating that these genes are likely direct targets of Foxn3 repression (Fig. 3C). An example is *Foxj1*, which had a prominent Foxn3 binding peak present at its promoter (Fig. 3D). The Foxn3 protein has been shown to bind preferentially to two different, unrelated DNA sequence motifs, referred to as FHL-N, and FKH (Nakagawa et al., 2013; Rogers et al., 2019). The FHL-N motif is a subset of the FHL binding motif bound by many members of the Forkhead family (Nakagawa et al., 2013). FHL-N, FHL, and FKH motifs were all significantly enriched in Foxn3 binding regions (Fig. 3E). Intriguingly, the most enriched de novo motif in the Foxn3 binding regions was similar to motifs bound by RFX family transcription factors, which have been linked to cilia gene regulation (Choksi et al., 2014). Motifs for both Rfx3 (a dimeric RFX binding motif) and Rfx7 (a monomeric RFX binding motif) were enriched in regions bound by Foxn3 (Fig. 3E).

**Figure 3.**
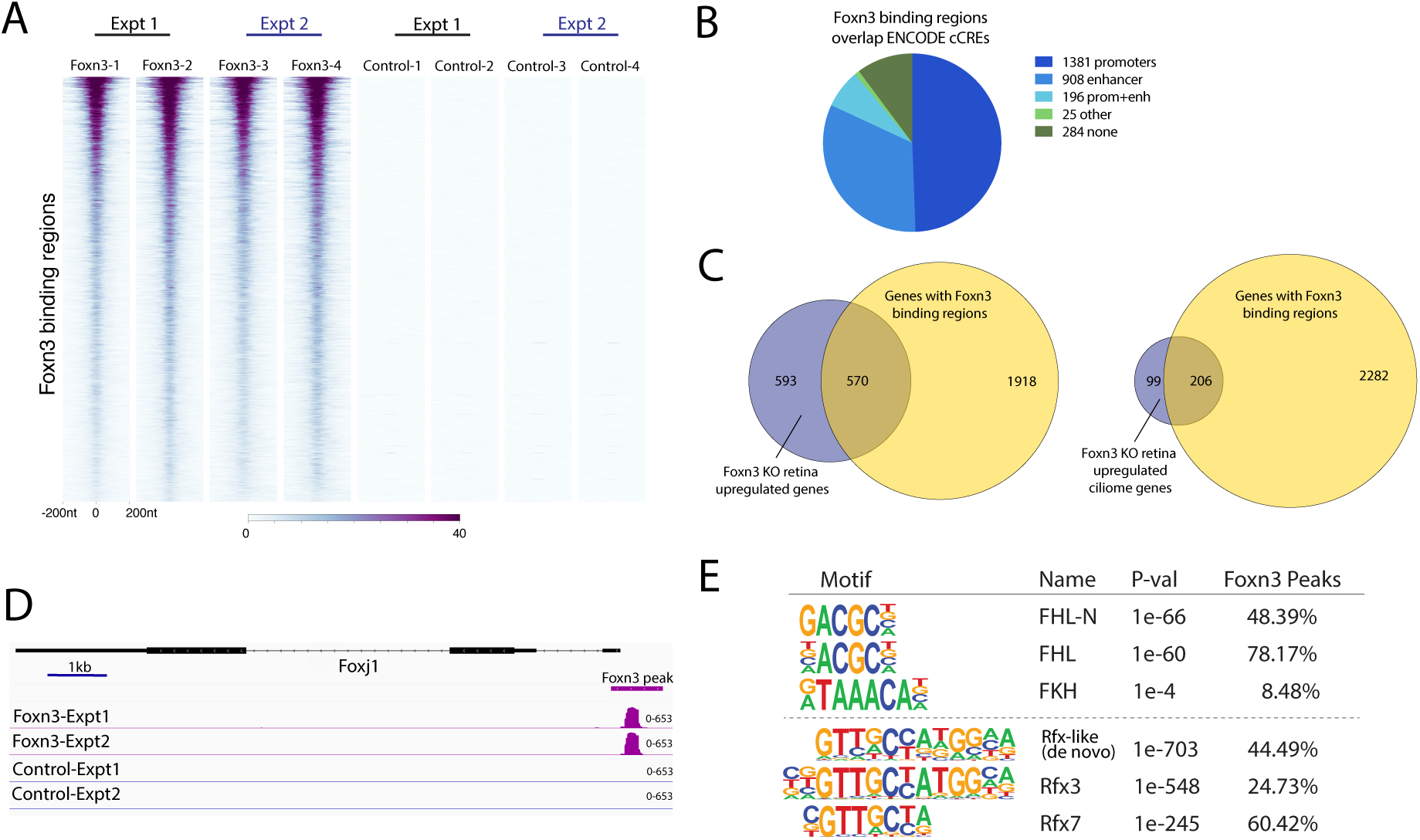
Analysis of Foxn3 genomic binding in the developing retina. (A) Heat map of read coverage at consistent Foxn3 binding regions identified by CUTCRUN in E16 retinas. Two anti-Foxn3 samples and two IgG control samples are shown for each of two experiments. (B) Most Foxn3 binding regions overlap Encode cCREs annotated as promoters or enhancers. (C) Venn diagrams of overlap between significantly upregulated genes or significantly upregulated ciliome genes in *Foxn3* KO retinas and genes with Foxn3 binding sites identified by CUTCRUN. (D) Foxn3 CUTCRUN read peaks at the *Foxj1* promoter. The range of read counts is indicated. (E) FHL-N, FHL, and FKH forkhead binding motifs are enriched in the Foxn3 binding regions. In addition, motifs for RFX transcription factors are enriched in the Foxn3 binding regions.

Prominent Foxn3 binding peaks were present at the promoters of *Ccdc39*, *Tekt1*, and *Cfap206* (Fig. 3A), three genes involved in cilia formation that are upregulated in Foxn3 KO retinas (Fig. 2A). We constructed Luciferase reporters driven by the promoter region from each gene and co-transfected HEK293 cells with the reporters and expression vectors for Foxj1, Rfx3, and/or Foxn3. We also tested whether the Rfx3 protein could activate the reporters and/or cooperate with Foxj1. Foxj1 activated all three reporters ∼8-15 fold (Fig. 4B-D), while Rfx3 by itself activated the reporters less than two-fold, but modestly increased Ccdc39 reporter activation when co-expressed with Foxj1. Co-expression of Foxn3 blocked activation of all three reporters by Foxj1, with or without Rfx3, consistent with Foxn3 directly repressing these genes (Fig. 4B-D).

**Figure 4.**
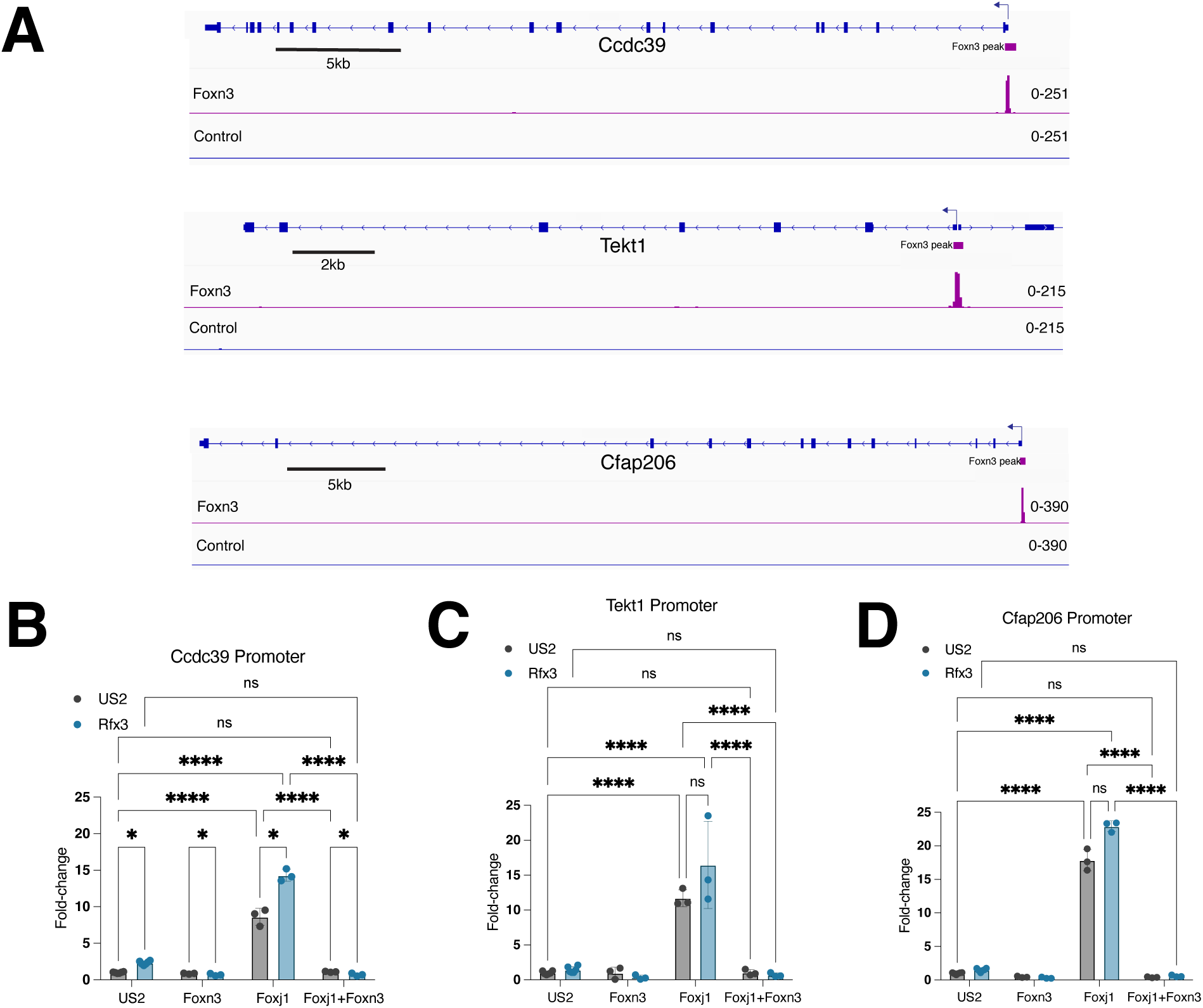
Foxn3 represses reporters for cilia genes. (A) CUTCRUN read coverage for three genes encoding proteins that function in primary cilia. Foxn3 binding regions are present at the promoters in E16 retinas. The range for read coverage for each sample is indicated. Graphs showing activation of transfected Luciferase reporters, based on the promoter regions from *Ccdc3S* (B), *Tekt1* (C), or *Cfap20C* (D), by cotransfection of a Foxj1 expression vector in HEK293 cells, relative to the US2 control vector, with or without cotransfection of either an Rfx3 expression vector or the US2 control vector. Reporter activation was by Foxj1 was completely blocked by co-expression of Foxn3. N=3-6, with standard deviations indicated. 2-way ANOVA: **** p≤0.0001, *p≤0.05.

To investigate whether Foxn3 genomic binding overlaps with genomic binding for the Rfx3 or Foxn4 proteins in the retina, we used CUTCRUN to identify regions bound by endogenous Rfx3 or Foxn4 in WT E16 mouse retinas (Fig. 5A-B) (Supplemental Tables 3, 4). Individual CUTCRUN samples showed consistent enrichment at the called peaks (Supplemental Fig. S4). Remarkably, 81% of the Foxn3 binding regions overlapped with regions bound by Rfx3, while 40% of the Foxn3 binding regions overlapped with regions bound by Foxn4; 32% of the Foxn3 binding regions overlapped with both Foxn4 and Rfx3 bound regions (Fig. 5C). For Foxn4, 36% of the bound regions overlapped with Rfx3 bound regions. Rfx3 binding motifs were significantly enriched in the regions bound by Rfx3, while FHL-N and FHL binding motifs were significantly enriched in the Foxn4 bound regions (Fig. 5D).

**Figure 5.**
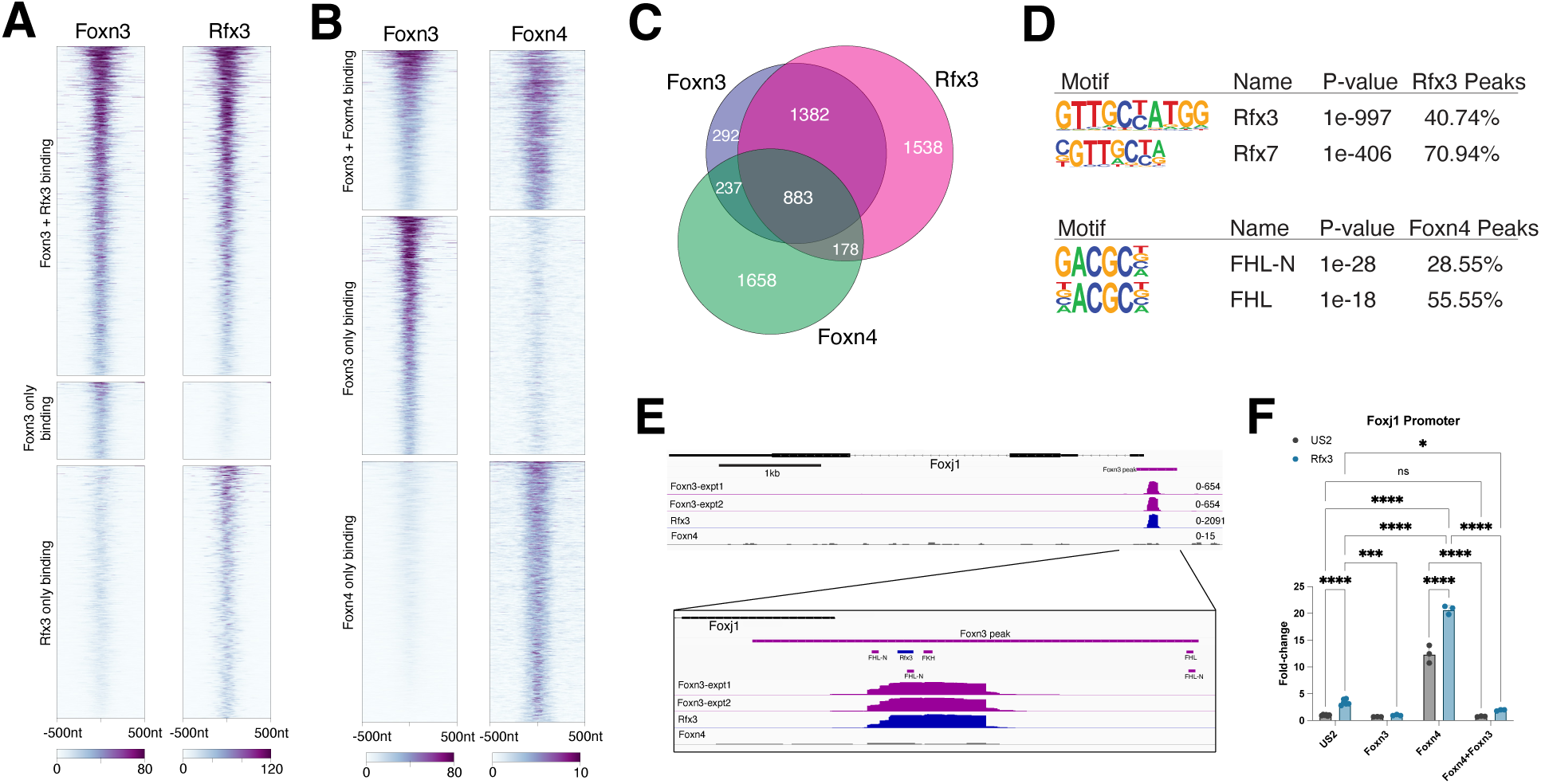
Analysis of Rfx3 and Foxn4 genomic binding in the developing retina. (A) Heat map showing Rfx3 binding regions identified by CUTCRUN. Most Foxn3 binding regions overlap with Rfx3 binding regions. (B) Heat map showing Foxn4 binding regions identified by CUTCRUN. Many Foxn4 binding regions overlap with Foxn3 binding regions. Heat maps for individual samples are shown in Supplemental Figure 4. (C) Venn diagram showing overlaps among Foxn3, Rfx3, and Foxn4 binding regions. (D) Rfx3 (dimeric) and Rfx7 (monomeric) binding motifs are enriched in the Rfx3 bound regions. FHL-N and FHL motifs are enriched in the Foxn4 bound regions. (E) In E16 retinas, the *Foxj1* promoter region is bound by Foxn3 and by Rfx3, based on CUTCRUN, but not by Foxn4. Locations of Rfx3 or FHL/FKH motifs and read counts are indicated. (F) Luciferase reporter based on *Foxj1* promoter was cooperatively activated by Foxn4 and Rfx3 in HEK293 cells, but co-expression of Foxn3 prevented activation by Foxn4 with or without Rfx3, and reduced activation by Rfx3 alone. N=3-6 with standard deviations indicated. 2-way ANOVA: **** p≤0.0001, *** p≤0.001, *p≤0.05, ns= not significant.

The *Foxj1* promoter region was bound by both Foxn3 and Rfx3 in the embryonic retinas, but not by Foxn4 (Fig. 5E). We constructed a Luciferase reporter with the *Foxj1* promoter region and co-transfected it into HEK293 cells with expression vectors for Foxn4, Rfx3, and/or Foxn3 (Fig. 5F). Rfx3 weakly activated the Foxj1 promoter reporter, while Foxn4 activated more than 10-fold, and co-expression of Rfx3 with Foxn4 further increased activation. However, co-expression of Foxn3 strongly inhibited activation of the reporter by Foxn4, with or without Rfx3. We constructed additional reporters based on intronic enhancers in the *Ptprf*, *Meis1*, and *Notch1* genes, all which are bound by Foxn3 and Rfx3 in retinas, based on our CUTCRUN data (Supplemental Fig. S5A). The *Meis1* and *Notch1* genes have been linked to the *Foxn4* pathway in retina and/or spinal cord (Islam et al., 2013; Luo et al., 2012; Misra et al., 2014), and both enhancers were bound by Foxn4 in the retina, while the *Ptrpf* enhancer was not. However, all three reporters could be activated by Foxn4 in HEK293 cells, and the *Ptprf* and *Meis1* reporters were further activated by co-expression of Rfx3 with Foxn4 (Supplemental Fig. S5B-D). While *Ptprf* was upregulated in the *Foxn3* KO retinas, neither Meis1 nor Notch1 mRNAs changed significantly (Supplemental Table 1). Surprisingly, only the *Ptprf* reporter was repressed by co-expression of Foxn3 (Supplemental Fig. S5B-D), suggesting that additional elements within the enhancers can modulate Foxn3 repression.

Luciferase reporters based on two intronic enhancers located in the *Foxn3* gene, each bound by Foxn3, Rfx3, and Foxn4, were constructed and assayed in HEK293 cells (Fig. 6A-E, Supplemental Fig. S5A, E). Both reporters were activated by Foxn4 and strongly activated by Foxn4 in combination with Ascl1, a basic-helix-loop-helix (bHLH) transcription factor that cooperates with Foxn4 in the retina (Luo et al., 2012) (Fig. 6B, Supplemental Fig. S5E). Co-expression of Rfx3 with either Foxn4 or with Foxn4 and Ascl1 together further increased activation of the reporters, while co-expression of Foxn3 strongly inhibited reporter activity driven by Foxn4, Ascl1, or Rfx3, alone or in combination (Fig. 6B, Supplemental Fig. S5E). For the *Foxn3* enhancer 2 reporter, we tested whether binding sites for each factor (Fig. 6A) were required for reporter function. Altering the two E-boxes (bHLH binding motifs) present in the reporter prevented Ascl1 cooperation with Foxn4 and Rfx3 (Fig. 6C). Mutation of the FKH and the FHL-N forkhead motifs together in the Foxn3 Enhancer 2 reporter greatly reduced activation by Foxn4 or by Rfx3 with Foxn4 (Fig. 6D). However, residual activation by Foxn4 still was strongly inhibited by co-expression of Foxn3 for either the reporter with E-box mutations or the reporter with FKH/FHL-N mutations (Fig. 6C, D). Mutation of the single Rfx3 binding motif present in the *Foxn3* enhancer 2 reporter greatly reduced activation by Foxn4 with Ascl1 (Fig. 6E). Furthermore, co-expression of Foxn3 failed to inhibit the reporter without the Rfx3 site when activated by Foxn4 and Ascl1. Taken together, these observations indicate that Rfx3 cooperates with Foxn4 for transcriptional activation and with Foxn3 for transcriptional repression.

**Figure 6.**
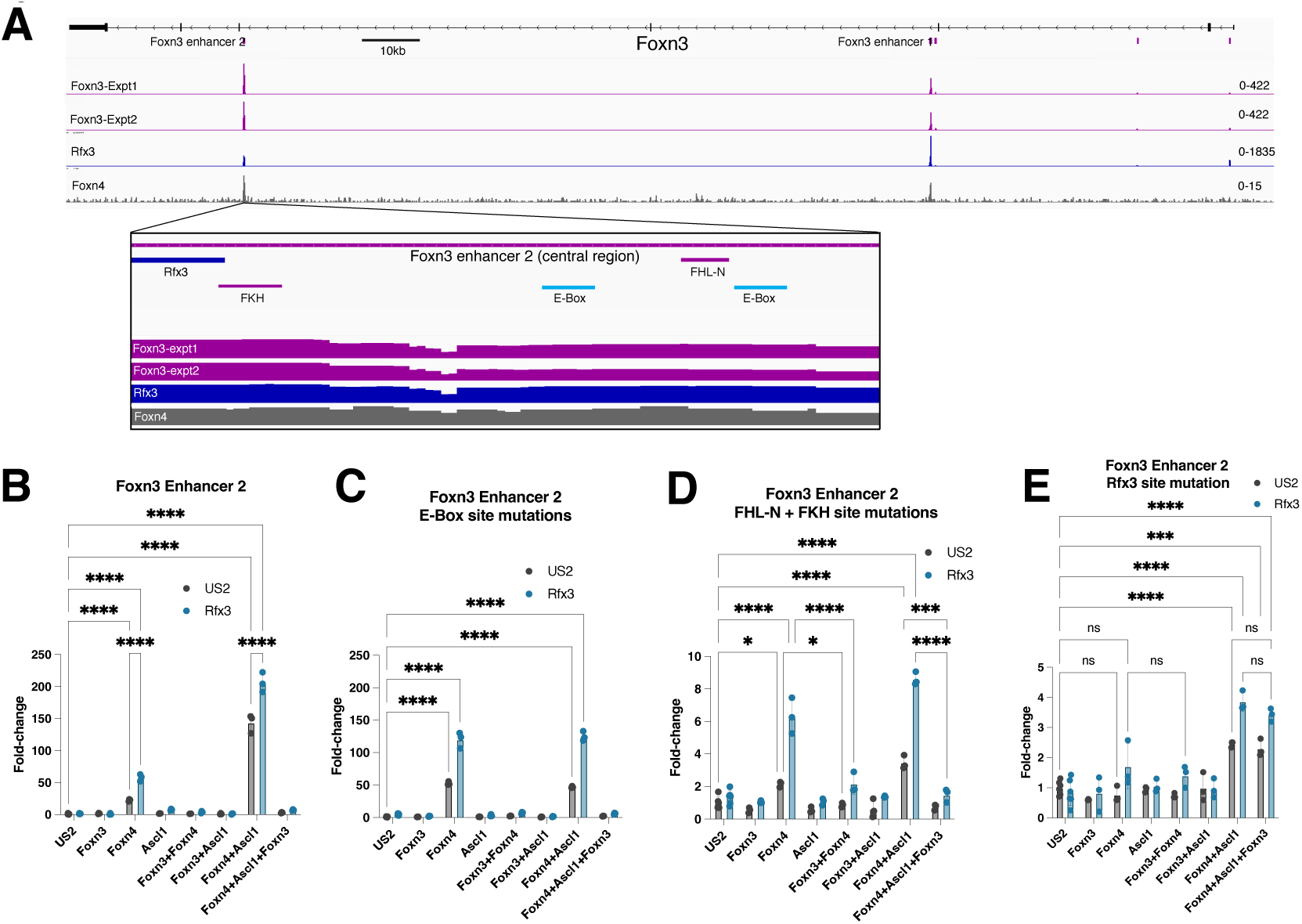
Regulation of a reporter by Foxn4 or Foxn3 requires an Rfx3 binding site. (A) CUTCRUN detects binding by Foxn3, Rfx3 and Foxn4 at a *Foxn3* enhancer. Read counts and potential binding site motifs for Rfx3, Ascl1 (E-boxes), and Foxn3/Foxn4 (FKH, FHL-N) are indicated. (B) A reporter based on the central region of a *Foxn3* enhancer was cooperatively activated by Foxn4, Ascl1, and Rfx3 and efficiently repressed by co-expression of Foxn3 in transfection assays in HEK293 cells. (C-E). Reporters with mutations in specific binding motifs were assayed. Disruption of the two E-boxes (C) prevented Ascl1 cooperation with Foxn4 but did not alter regulation by Foxn4 or Foxn3. Mutation of the FKH and FHL-N sites (D) greatly reduced but did not abolish activation by Foxn4. Residual activation was still repressed by Foxn3, with or without Rfx3. Mutation of the Rfx3 site (E) abolished or greatly reduced reporter activation by Foxn4 with or without Ascl1 and Rfx3, while Foxn3 failed to repress the remaining activation after mutation of the Rfx3 site. N=3-6. Labels as in Fig. 5.

### A conserved LXXLXWL motif in Foxn4, Foxj1, and Foxn3 is required for interaction with Rfx3

The mouse Foxn4 or Foxj1 proteins can functionally substitute for the C. elegans FKH-8 forkhead protein to support ciliome gene expression in C. elegans (Brocal-Ruiz et al., 2023). FKH-8 cooperates with DAF-19, the only RFX protein present in C. elegans, to activate cilia gene expression, while the Foxj1 protein interacts with RFX proteins to cooperatively regulate genes involved in motile cilia formation in mammals (Didon et al., 2013). Protein-protein interaction screens have detected associations between Foxj1, Foxj2, Foxn2, Foxn3 or Foxn4 and Rfx1, Rfx2 or Rfx3 (Huttlin et al., 2021; Huttlin et al., 2017; Li et al., 2015). We looked for amino acid sequences outside of the forkhead domain that are conserved among FKH-8, Foxj1, and Foxn2/Foxn3/Foxn4 to identify candidate motifs that might mediate complex formation with RFX proteins. A hydrophobic, leucine rich motif with a conserved tryptophan, LXXLXWL, is located N-terminal to the forkhead DNA binding region in these proteins (Fig. 7A). A similar sequence with the central leucine substituted by isoleucine or methionine is present in Foxj2 and Foxj3. A divergent motif without the central hydrophobic residue or the conserved tryptophan is present in mouse Foxi1, a protein that cannot substitute for FKH-8 in C. elegans (Brocal-Ruiz et al., 2023). We constructed expression vectors for Foxn4, Foxn3, and Foxj1 in which the LXXLXWL sequence was replaced with seven alanine residues (Foxn4-7A, Foxn3-7A, and Foxj1-7A, Fig. 7A). The Foxn3-7A protein remains capable of binding to DNA in an electromobility shift assay (EMSA), indicating that the 7A substitution does not disrupt the overall structure or expression of the protein (Supplemental Fig. S6). Hemagglutinin-(HA-) epitope-tagged versions of the Foxn4, Foxn3, or Foxj1 proteins co-immunoprecipitated with Rfx3 when the co-expressed in HEK293 cells, but HA-tagged versions of the Foxn4-7A, Foxn3-7A, and Foxj1-7A proteins did not (Fig. 7B). Foxn4-7A failed to activate the *Foxn3* enhancer 2 reporter by itself and showed reduced activation when combined with Rfx3. Foxn4-7A had reduced reporter activation in combination with Ascl1 and Rfx3. The Foxn3-7A expression vector failed to inhibit activation of the reporter by Foxn4, with or without co-expression of Ascl1 and Rfx3 (Fig. 7C). Together with previous reports, these observations indicate that the Rfx3 protein forms a complex with Foxn3, Foxn4, or Foxj1 to promote transcriptional regulation of target genes, and that all three forkhead proteins require the conserved LXXLXWL motif for complex formation with Rfx3.

**Figure 7.**
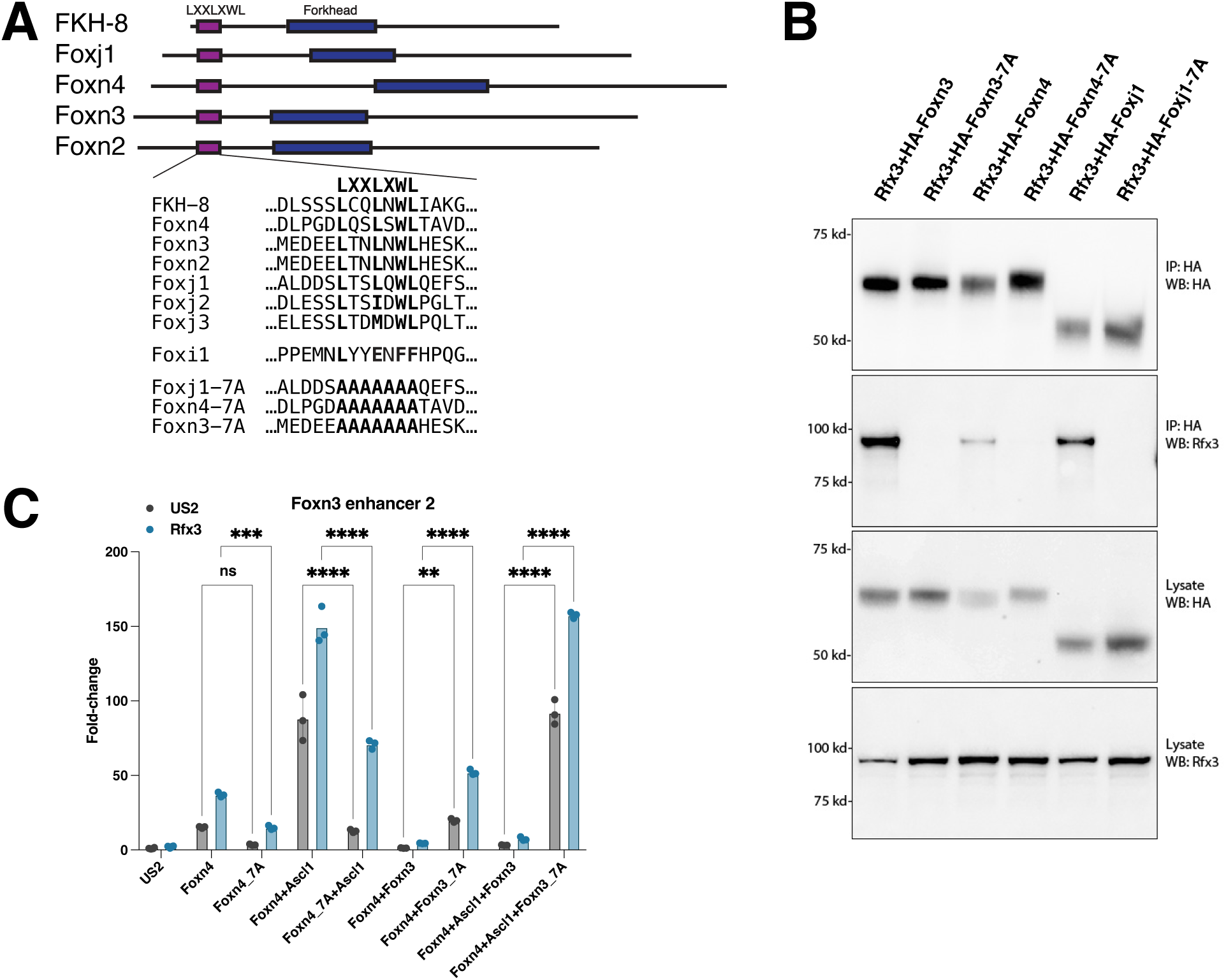
A conserved protein motif required for complex formation and cooperation between Rfx3 and forkhead factors. (A) Schematic showing the location of an LXXLXWL motif shared by the C. elegans FKH-8 protein and several mouse forkhead transcription factors, including Foxn3, Foxn4, and Foxj1. (B) Epitope tagged HA-Foxn4, HA-Foxj1, or HA-Foxn3 co-immunoprecipitated (IP) with Rfx3 when co-expressed in HEK293 cells. Substitution of the LXXLXWL motif with 7 alanines in HA-Foxn3-7A, HA-Foxn4-7A, or HA-Foxj1-7A (A) prevented co-immunoprecipitation with Rfx3. WB: representative Western blot (N=3). (C) Foxn4-7A had reduced reporter activation compared to Foxn4, with or without Rfx3. In contrast to Foxn3, Foxn3-7A failed to repress activation of the reporter by Foxn4 by itself or in combination with other factors. N=3-6, 2-way ANOVA: ** p<0.01. Other labels as in Fig. 5. See Supplemental Figure 7 for complete western blots.

### Interaction between the LXXLXWL motif and the Rfx3 dimerization domain

Complete structures are not available for the Foxn3 or Rfx3 proteins, although structures for the RFX1 and Foxn3 DNA binding domains have been reported (Gajiwala et al., 2000; Rogers et al., 2019). We used AlphaFold 3 (Abramson et al., 2024) to model the structure of Foxn3 in a complex with a dimer of two Rfx3 proteins and a short DNA sequence with Rfx3 and FKH sites. Interestingly, the predicted structure juxtaposed the Foxn3 LXXLXWL motif to the interface between the two Rfx3 dimerization domains, with the LXXLXWL motif as the main contact site for Foxn3 with the two Rfx3 proteins (Supplemental Fig. S8A-C). PyMol indicates a pi-pi interaction between the tryptophan (W65) in the Foxn3 LXXLXWL motif and a conserved phenylalanine (F620) in one Rfx3 molecule of the predicted structure. However, overall confidence scores for the predicted complex were low, likely due to substantial disordered regions present in each protein. We therefore used AlphaFold 3 to generate a model based on the C-terminus of Rfx3, including the extended dimerization domain and the adjacent B domain (Emery et al., 1996; Katan-Khaykovich and Shaul, 1998), in combination with the N-terminus of Foxn3 (Fig. 8A-B, Supplemental Fig. S9). This model placed the Foxn3 LXXLXWL motif at the same interface with the paired Rfx3 dimerization domains and maintained the W65 pi-pi interaction with Rfx3 F620, but with high confidence scores (Supplemental Fig. S8C). AlphaFold 3 models for the N-terminus of Foxn4 or Foxj1 with the Rfx3 dimerization domain yielded similar high confidence score predictions for interaction via the LXXLXWL motif, also with a pi-pi interaction between the tryptophan and Rfx3 F620 (Supplemental Figs. S10 and S8C). In Foxj1 a second tryptophan (W48) located near the LXXLXWL motif, is also predicted to form a pi-pi interaction with Rfx3 F620.

**Figure 8.**
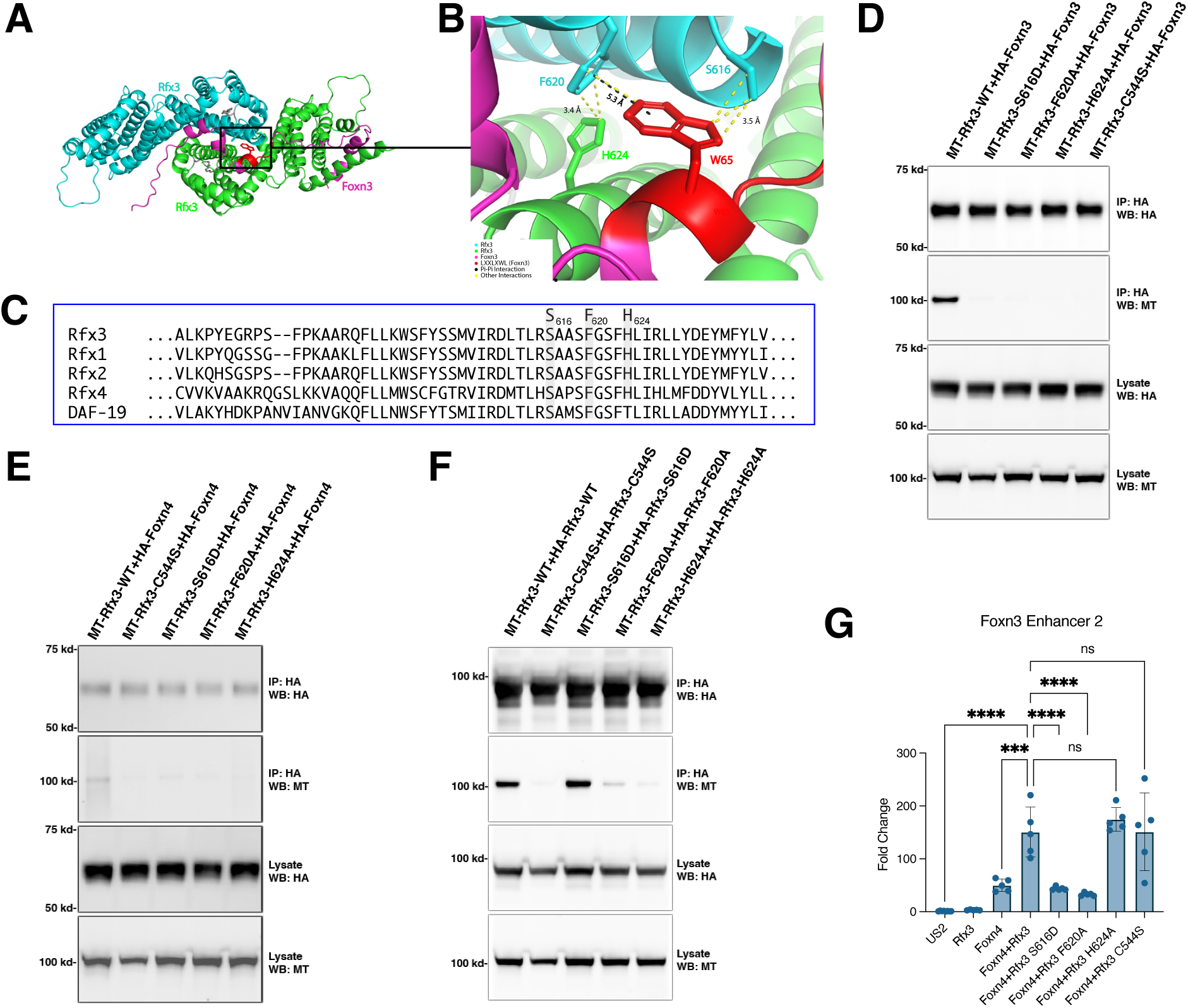
Mutations within the Rfx3 dimerization domain disrupt interaction and cooperation with forkhead proteins. (A, B) Interaction of the Foxn3 N-terminus with a dimer of the Rfx3 C-terminus modeled by AlphaFold 3. Rfx3 molecules are palmitoylated at C544 (gray). A) Full view of Rfx3 dimerization domain (two molecules) and Foxn3 N-terminus interaction with each protein labelled. The Foxn3 LXXLXWL motif (red) contacts the dimerization interface between the two Rfx3 proteins. (B) Enlarged image of the interaction site. W65 in the Foxn3 LXXLXWL motif is labelled along with S616, F620, and H624 residues from Rfx3. Predicted pi-pi interaction between F620 of Rfx3 and W65 of Foxn3 (distance: 5.3 Å); other interactions between the four amino acids indicated. C) Alignment of predicted forkhead interaction region in the mouse Rfx3 dimerization domain with other mouse RFX proteins and the C. elegans DAF-19 RFX protein, with the 3 amino acids selected for mutation indicated. (D) Epitope tagged MT-Rfx3 and HA-Foxn3 co-immunoprecipitated when co-expressed in HEK293 cells but HA-Foxn3 and MT-Rfx3 with S616D, F620A, H624A, or C544S mutations did not. (E) HA-Foxn4 also co-immunoprecipitated with MT-Rfx3 but not with MT-Rfx3 containing S616D, F620A, H624A, or C544S mutations. (F) Co-immunoprecipitation of MT-Rfx3 and HA-Rfx3 shows that WT Rfx3 or Rfx3 S616D can homodimerize, but the F620A, H624A, or C544S mutations reduce homodimerization. (G) Co-expressed Rfx3, Rfx3 H624A, or Rfx3 C544S cooperated with Foxn4 to activate the *Foxn3* Enhancer 2 luciferase reporter, while Rfx3 S616D or Rfx3 F620A failed to cooperatively activate with Foxn4. N=5-8. One-way ANOVA: **** p≤0.0001, *** p≤0.001, ns= not significant. See Supplemental Figures 8-13 for related data and complete western blots.

To investigate amino acids in Rfx3 required for interaction with forkhead proteins, we generated expression vectors for Rfx3 with individual substitution mutations: F620A, S616D, or H624A. Rfx3 S616 is predicted to interact with the Foxn3 LXXLXWL motif, while Rfx3 H624 is predicted to interact with Rfx3 F620 in the other chain of the Rfx3 dimer (Fig. 8B). F620 and S616 are conserved among Rfx1-4 and DAF-19 (Fig. 8C). HA-tagged Foxn3 or HA-tagged Foxn4 were co-expressed with Rfx3 or each of the mutated Rfx3 proteins in HEK293 cells. In addition, we tested Rfx3 C544S, a mutation that disrupts Rfx3 dimerization (Chen et al., 2018) but which is not located near the predicted interaction with the LXXLXWL motif (Supplemental Figure S8D). Rfx3 proteins with each mutation showed little co-immunoprecipitation with either Foxn3 or Foxn4 (Fig. 8D). We evaluated the effect of each mutation on Rfx3 dimerization by co-immunoprecipitation of HA- and Myc-tagged Rfx3 proteins with the same mutation. The F620A, H624A, and C544S substitutions reduced co-immunoprecipitation of Rfx3 dimers, while the S616D substitution did not (Fig. 8E). Consistent with these results, F620 and H624 are predicted to be located at the Rfx3 dimerization interface and interact with residues in the other Rfx3 chain, while Rfx3 S616 appears to interact only with the LXXLXWL motif (Fig. 8A-B). The S616D, F620A, and H624A mutations in Rfx3 did not prevent DNA binding as assessed by EMSA (Supplemental Fig. S14). However, only Rfx3 S616D showed the same pattern of shifted bands as WT Rfx3. The altered EMSA patterns for Rfx3 F620A or Rfx3 H624A resemble EMSA changes reported for Rfx1 proteins that cannot dimerize (Katan-Khaykovich and Shaul, 1998), consistent with the reduced dimerization observed for Rfx3 F620A and Rfx3 H624A. Co-expression of Foxn4 with either Rfx3 S616D or Rfx3 F620A reduced cooperative activation of a reporter, while co-expression of Foxn4 with Rfx3 H624A or Rfx3 C544S fully activated the reporter (Fig.8F). Taken together, these observations support a direct interaction between the forkhead LXXLXWL motif and the dimerization domains of an Rfx3 dimer. While disruption of Rfx3 dimerization prevented co-immunoprecipitation with Foxn3 or Foxn4, only mutations at Rfx3 S616 or Rfx3 F620, two residues predicted to interact directly with the forkhead LXXLXWL motif, prevented cooperative reporter activation with Foxn4.

### A potential truncated Foxn3 protein in Foxn3 KO mice does not regulate a reporter

*Foxn3* KO mice express a Foxn3 mRNA in which exon 2 splices to exon 4 based on RNA-seq. The N-terminus of Foxn3 is present in exon 2 and includes the LXXLXWL motif and part of the forkhead domain. The C-terminus of Foxn3, encoded by exons 4 and 5, is out of frame when exon 2 splices to exon 4, potentially expressing a truncated Foxn3 protein with a novel C-terminus in *Foxn3* KO mice. Cotransfection of an expression vector for the Foxn3 coding region without exon 3 had no effect on the activation of a reporter by Foxn4, Ascl1, and Rfx3 in HEK293 cells, indicating that the potential truncated Foxn3 protein is likely not functional (Supplemental Fig. S15).

## Discussion

Over 300 genes involved in cilia formation or function were upregulated in the developing retinas of *Foxn3* KO mice, with some mRNAs increasing over 100-fold. CUTCRUN analysis revealed that most upregulated ciliome genes in the Foxn3 KO retinas were bound by the Foxn3 protein, consistent with direct repression of these genes by Foxn3 (Scott and Plon, 2005). In differentiated retinas, Foxn3 is expressed in bipolar cells and some amacrine cells (Zhang et al., 2022). Strikingly, loss of *Foxn3* led to increased cilia on bipolar cells and amacrine cells in the retina INL at P21, indicating that Foxn3 is required to prevent ectopic cilia formation in the mouse retina. The consequences of additional primary cilia on cellular function are unclear, but most retinal bipolar cells in the mouse do not have primary cilia (Ning et al., 2021), so bipolar cells with cilia potentially could respond to extracellular signals that might otherwise not be received. These results are consistent with the observation that overexpression of FOXN3 downregulated genes involved in cilium biogenesis and reduced primary cilia formation in human lung carcinoma cells (Huot et al., 2014).

We observed additional displaced amacrine cells in the GCL of *Foxn3* KO retinas at P21 as well as ectopic amacrine cells located within the IPL, but the number of amacrine cells, bipolar cells, and Müller glia in the INL were unchanged. Ectopic amacrine cells were not apparent at E19, suggesting that these amacrine cells are generated and/or migrate during postnatal retinal development. The amacrine cells located in the central IPL in *Foxn3* KO retinas resemble a layer of ectopic amacrine cells observed in the IPL of *Plagl1* (*Zac1*) KO retina explants (Ma et al., 2007), although there are fewer ectopic amacrine cells in the IPL of *Foxn3* KO retinas. The loss of *Plagl1* reduces TGF-beta II feedback inhibition of amacrine cells. Altered TGF-beta signaling could occur in the Foxn3 KO retinas since TGF-beta signaling is mediated by receptors in primary cilia and signaling pathways can be impacted by changes within cilia (Christensen et al., 2017). However, a recent report linked reduced Foxn3 to the stabilization of Smad4 and increased TGF-beta signaling in lungs (Yu et al., 2025), suggesting that Foxn3 could regulate the pathway at multiple levels.

*Foxn3* KO retinas had additional retinal astrocytes in the GCL and IPL, with astrocytic processes extending throughout the IPL. Astrocytes migrate into the retina from the optic nerve head (reviewed by Paisley and Kay, 2021) and their dispersal across the retina is guided by signals from retinal ganglion cells (O’Sullivan et al., 2017). Hypoxia can trigger proliferation of astrocytes in neonatal mouse retinas (Perelli et al., 2021). Transgenic overexpression of PDGF-A in retinal neurons also increases the number of astrocytes and leads to spread of astrocytes into additional layers of the retina (Fruttiger et al., 1996), suggesting that the changes in the *Foxn3* KO retinas could arise from altered signaling in primary cilia on astrocytes. Consistent with this possibility, the maturation and function of brain astrocytes depend on primary cilia during development (Wang et al., 2024). However, Foxn3 also regulates genes that are not associated with cilia, so it remains possible that developmental changes in astrocytes and/or amacrine cells arise from gene expression changes unrelated to cilia.

While this manuscript was in preparation, Liu et al. reported that retina-specific conditional knockout (cKO) of *Foxn3* led to upregulation of cilia-associated genes, including Foxj1, and the formation of ectopic cilia on bipolar and amacrine neurons in mature retinas (Liu et al., 2025), consistent with the results described here for *Foxn3* KO retinas. Retinal astrocytes were not analyzed in the *Foxn3* cKO, and additional displaced amacrine cells or IPL-localized amacrine cells were not reported in the *Foxn3* cKO retinas. This difference may arise in part from the models used: in the *Foxn3* cKO, *Foxn3* exon 2 was deleted in the developing retina with *Six3-Cre*, while in the model analyzed here, *Foxn3* exon3 was deleted in all cells throughout embryogenesis. In addition, retinas from the *Foxn3* cKO were analyzed at later ages, so potential ectopic amacrine cells at P21 may have been reduced or eliminated by cell death.

Forkhead proteins and RFX transcription factors have been linked to the activation of cilia gene expression in both vertebrates and invertebrates, including Foxj1 and Foxn4. The promoter of *Foxj1* was bound by Foxn3 in embryonic retinas, and the Foxj1 mRNA was upregulated more than 20-fold in *Foxn3* KO retinas. Increased Foxj1 protein expression likely further activates Foxj1 targets derepressed by the absence of Foxn3. Foxj1 cooperates with Rfx3 and we found that Rfx3 genomic binding in the embryonic mouse retina extensively overlaps with Foxn3 binding and overlaps to a lesser extent with Foxn4 binding. Reporters based on genomic regulatory elements bound by these proteins could be cooperatively activated by Foxn4 and Rfx3 in cell culture but were repressed by Foxn3. While Rfx3 expression by itself had modest effects on reporters, mutation of the Rfx3 binding site in a reporter both reduced cooperative activation by Foxn4 and prevented repression by Foxn3, indicating that different forkhead proteins can cooperate with Rfx3 to either activate or repress target genes.

Interactions between RFX proteins and the Foxj1, Foxn4, Foxn2, and Foxn3 forkhead proteins have been reported (Didon et al., 2013; Huttlin et al., 2021; Huttlin et al., 2017; Li et al., 2015). We identified an evolutionarily conserved LXXLXWL motif present in the C. elegans FKH-8 and mouse Foxj1, Foxn4, Foxn2, and Foxn3 proteins that is required for association between Rfx3 and the Foxj1, Foxn4, or Foxn3 proteins. Mutation of the LXXLXWL motif in Foxn4 or Foxn3 reduced cooperative activation of a reporter by Foxn4 with Rfx3 and prevented Foxn3 repression of the reporter. AlphaFold 3 predicted that the forkhead LXXLXWL motifs in Foxn3, Foxn4 or Foxj1 interact with the Rfx3 dimerization domain in an Rfx3 homodimer. Single amino acid substitutions in Rfx3 at the predicted interface with the LXXLXWL motif, or mutations that disrupt Rfx3 homodimer formation, prevented co-immunoprecipitation of Rfx3 with Foxn3 or Foxn4, supporting the predicted interaction. In contrast, only substitutions at either of two amino acids in Rfx3 that are predicted to directly interact with the LXXLXWL motif prevented Rfx3 from cooperating with Foxn4 on a reporter. Mutations that disrupted Rfx3 dimerization altered but did not prevent Rfx3 DNA binding as assessed by EMSA, consistent with prior analysis of Rfx1 (Katan-Khaykovich and Shaul, 1998). Interaction with DNA and/or other proteins may be sufficient to allow the formation of Rfx3 dimers on DNA even when Rfx3 cannot dimerize in solution. Alternately, an Rfx3 monomer may interact with forkhead proteins when bound to adjacent sites on DNA. Based on our observations, cooperation between Foxn4 or Foxj1 and Rfx3 likely involves both binding to nearby genomic sites and the formation of protein-protein complexes. The Foxn3 protein includes transcriptional repression domains located C-terminal to the forkhead domain (Huot et al., 2014; Scott and Plon, 2005). Since Foxn3 can bind at the same sequences as Foxn4 or Foxj1 (Nakagawa et al., 2013; Rogers et al., 2019), Foxn3 may repress genes by competing with Foxn4 or Foxj1 for DNA binding sites, by competing for interaction with Rfx3, and by recruiting transcriptional corepressors. Such a multifaceted mechanism should allow more potent repression by Foxn3 (see model in Supplemental Figure S16). As the Rfx3 residues required for interaction with the LXXLXWL motif are conserved in other RFX factors (Fig. 8C), additional RFX proteins likely interact with forkhead factors that contain the LXXLXWL motif.

RFX proteins have been associated with both activation and repression of target genes (Didon et al., 2013; Iwama et al., 1999; Nakayama et al., 2003). The extensive overlap between Rfx3 and Foxn3 genomic binding in the developing retina raises the possibility that Foxn3 represses numerous genes in part by interaction with Rfx3. Some of the shared binding sites for Foxn3 and Rfx3 detected by CUTCRUN contain one or more RFX binding motifs but do not contain a consensus Foxn3 binding motif (e.g., see *Ptprf* in Supplemental Fig. S5A). These observations suggest that Foxn3 could be recruited into complexes with DNA-bound Rfx3 without direct DNA binding by Foxn3. Alternately, Foxn3 in a complex with Rfx3 may be able to bind to a non-canonical DNA binding site, as reported for cooperative binding by other transcription factors (Berkes et al., 2004). Genetic disruptions of Rfx3 and other RFX factors are associated with human neurodevelopmental disorders (Harris et al., 2021), raising the possibility that interactions between RFX factors and Foxn3 or other forkhead factors could play important roles in CNS development.

We previously found that Foxn3 and Foxn4 mRNAs expression overlaps in retinal progenitor cells (Zhang et al., 2022). Since Foxn4 binds to two intronic enhancers in the *Foxn3* gene, and can activate reporters based on those enhancers, it is possible that Foxn4, in cooperation with Ascl1 and/or other factors, upregulates Foxn3 in retinal cells to prevent inappropriate activation of the *Foxj1* gene and its downstream targets in the motile cilia pathway. However, Foxn3 mRNA increased about two-fold in Foxn4 KO retinas (see Supplemental Table 2 in Liu et al., 2020), suggesting that other factors also influence Foxn3 expression. Foxn3 mRNA also increases in the *Foxn3* KO retinas, and Foxn3 appears likely to repress its own expression based on its ability to repress reporters based on two *Foxn3* intronic enhancers, suggesting negative feedback may limit Foxn3 expression.

We identified the Foxn3 mRNA as a target of miR-216b (Zhang et al., 2022). Overexpression of miR-216a/b or inhibition of Foxn3 expression in neonatal retinas led to the formation of additional amacrine interneurons and loss of bipolar interneurons. Here we find that disruption of Foxn3 alters amacrine cells localization and number in the postnatal retina, although effects from the Foxn3 KO differ from miR-216a/b overexpression. Most additional amacrine cells observed after miR-216a/b overexpression were in the INL, but we did observe additional displaced amacrine cells (Zhang et al., 2022). Since *Foxn3* is disrupted throughout development in *Foxn3* KO mice, it is possible that some compensation occurs in these mice. Alternately, miR-216a/b may regulate additional targets that contribute to the observed phenotypes. We proposed a model in which inhibition of Foxn3 expression by miR-216a/b allowed increased Foxn4 activity to promote amacrine cell formation. However, the results presented here suggest that miR-216a/b and Foxn3 could affect amacrine cell development through changes in primary cilia. Consistent with this, Argonaute PAR-CLIP identified a miR-216a/b site in the 3’ UTR of the Rfx3 mRNA in the P0 retina (see Supplemental Table 3 in Zhang et al. 2022), suggesting that miR-216a/b may target multiple regulators of cilia gene expression.

While our analysis focused on the role of Foxn3 in the retina, most *Foxn3* KO mice die at or shortly after birth and the few surviving mice are smaller than normal, indicating that *Foxn3* has additional important developmental functions. Craniofacial defects were reported in mice with a hypomorphic *Foxn3* allele and in frogs after morpholino inhibition of Foxn3 (Samaan et al., 2010; Schmidt et al., 2011; Schuff et al., 2007), but the *Foxn3* KO mice analyzed here do not show craniofacial defects. However, Foxn3 function has been implicated in the repression of cell growth, acting as a tumor suppressor, regulation of metabolism in liver, and regulation of the inflammatory response in lungs (Huot et al., 2014; Karanth et al., 2016; Li et al., 2017; Xue et al., 2019; Yu et al., 2025; Zhang et al., 2021; Zhu et al., 2023). The extensive regulation of genes involved in cilia by Foxn3 suggests that some of these roles may be linked to the proper formation or function of cilia.

## Methods

### Mice and processing of retinas

All mouse (*Mus musculus*) experiments were approved by the Institutional Animal Care & Use Committee at the University of Michigan. WT retinas for CUTCRUN analysis were isolated from embryonic day 16 (E16) embryos of CD-1 mice (Charles River). The C57BL/6NJ-*Foxn3^em1(IMPC)Bay^*/Mmnc mouse strain (RRID:MMRRC_071094-UNC), with a heterozygous deletion of Foxn3 exon 3, was obtained from the Mutant Mouse Resource and Research Center (MMRRC) at University of North Carolina at Chapel Hill, an NIH-funded strain repository, and was donated to the MMRRC by Jason Heaney, Ph.D., Baylor College of Medicine (Birling et al., 2021). C57BL/6NJ-*Foxn3^em1(IMPC)Bay^*/Mmnc mice were bred with WT C57/BL6J mice (The Jackson Laboratory). Heterozygous male and female offspring were interbred to generate Foxn3 KO embryos or offspring. The genotypes of offspring and embryos were determined by genomic PCR (see Supplemental Table 5 for primers).

For tissue analysis, eyes were fixed with 2% paraformaldehyde (ThermoFisher) in phosphate-buffered saline (PBS, Sigma) for 30 min at room temperature and retinas were dissected from the eyes. Retinas were cryoprotected in PBS containing 30% sucrose (Sigma) for 2 hrs at 4 °C, embedded in OCT (Sakura), and sectioned at 15 µm using a cryostat. For the latter, serial sections were collected at 300 µm intervals across the retina. All retinas were processed under identical conditions for antibody staining.

### Immunofluorescence Analysis

Antibodies used and dilutions are listed in Supplementary Table 5. For retinal sections, samples were blocked for 1 hr at room temperature in PBS containing 0.1% Triton X-100 (ThermoFisher), 1% unlabeled affinity-purified Fab fragment donkey anti-mouse IgG (H+L), and 5% donkey serum (Jackson ImmunoResearch). Sections were incubated overnight at 4 °C with primary antibodies: mouse anti-AP2α, rabbit anti-Sox9, goat anti-GFAP, sheep anti-Vsx2, mouse anti-Arl13b, mouse anti-Foxj1, and mouse anti-GS. The following day, sections were incubated for 1 hr at room temperature with secondary antibodies (1:1000, Jackson ImmunoResearch): Alexa Fluor 647–conjugated anti-mouse, Alexa Fluor 488–conjugated anti-goat, Alexa Fluor 488–conjugated anti-sheep, and Alexa Fluor 568–conjugated anti-rabbit. Nuclei were counterstained with Hoechst 33342 (Roche). Immunofluorescence was imaged using an Olympus FV3000 confocal microscope or Oxford Andor BC43 microscope, and images were analyzed with FV10-ASW 3.1 Viewer or Imaris Viewer 10.2.0. To assemble complete retina sections shown in Fig. 1 and Supplementary Fig. S1, overlapping images were stitched together using Oxford Instruments Andor Fusion 2.5.0.218.

For whole-mount preparations, fixed retinas were processed with blocking solution at room temperature for 2 hrs, followed by primary antibody incubation overnight at 4°C with mouse anti-Arl13b and rabbit anti-IFT88. Secondary antibody incubation (Alexa Fluor 647-conjugated anti-mouse and Alexa Fluor 568-conjugated anti-rabbit (1:1000) was performed for 2 hrs at room temperature. Hoechst 33342 was used to counterstain the nuclei. Retinas were flat-mounted in PBS, and immunofluorescence was imaged on an Oxford Andor BC43 microscope. Images were analyzed, and colors were assigned using Imaris Viewer 10.2.0 or FIJI.

For both sections and whole-mount preparations, representative images were cropped, assembled, and labeled using Adobe Photoshop and/or Adobe Illustrator.

### Cell Counts

For Fig.1, AP2α, Vsx2, and Sox9 INL cells were counted within a 200-µm region in the middle zone between the center and edge on each side of a stitched retina section. Sox9-positive cells in the IPL and GCL were quantified from complete stitched retina sections. For Fig.2, Arl13b-labeled cilia were quantified from a single 200-µm region on one side. Blinded counts were performed manually using the Photoshop count tool. Statistical analyses were performed with GraphPad Prism 10 using unpaired t-tests. The counting analyses included a total of 4–6 sections per retina, from three P21 Foxn3 KO mice and three P21 WT mice.

### Expression vectors and reporters

Plasmids were constructed using standard techniques. Oligonucleotide PCR primers for plasmid construction are in Supplemental Table 5. The pUS2, pUS2-MT, pUS2-Foxn3, and pUS2-Foxn4 plasmids have been previously described (Chung et al., 2006; Zhang et al., 2022). DNA oligonucleotides encoding two copies of the HA tag were inserted into pUS2, pUS2-Foxn3, or pUS2-Foxn4 between the BamHI and EcoRI sites, resulting in pUS2-HA, pUS2-HA-Foxn3, and pUS2-HA-Foxn4. The coding regions for mouse Foxj1, Rfx3, and Foxn3 without exon 3 (from C57BL/6NJ-*Foxn3^em1(IMPC)Bay^*/Mmnc mice) were isolated by RT-PCR, using RNA extracted from embryonic mouse retinas, and inserted into pUS2, pUS2-HA, or pUS2-MT. The 7A mutants, in which the LXXLXWL motif was replaced with seven alanines (LXXLXWL → AAAAAAA), and Rfx3 point mutants (C544S, S616D, F620A, and H624A) were generated by PCR-based site-directed mutagenesis.

Enhancer or promoter regions from *Foxn3*, *Tek1*, *Ccdc3S*, *Cfap20C*, *Notch1*, *Meis1*, *Foxj1*, and *Ptprf* were amplified from mouse genomic DNA via PCR and cloned into pGL4.29 (Promega) using the NEBuilder HiFi DNA Assembly Cloning Kit (New England Biolabs, NEB). The Foxn3_Enh2_FBM construct, which contains mutations in two forkhead binding sites, and the Foxn3_Enh2_EBM construct, with mutations in two E-boxes, were created by inserting synthesized oligonucleotides into pGL4.29: two DNA oligonucleotides with complementary 3′ ends were annealed and extended with Q5 DNA polymerase (NEB) to generate a double-stranded DNA fragment, which was then cloned into the plasmid. The Foxn3_Enh2_Rfx3M construct, with mutation of the Rfx3 binding site, was generated using PCR site-directed mutagenesis.

### RNA-seq

Retinas from individual genotyped E19 embryos (Foxn3 KO or WT) were isolated, and total RNA from each retina was purified using TRIzol (ThermoFisher). A total of 500 ng of RNA was used to prepare the RNA-seq library using the NEBNext Ultra™ II Directional RNA Library Prep Kit for Illumina and NEB Unique Dual Index UMI Adaptors (NEB), following the manufacturer’s instructions. The RNA-seq libraries were sequenced on an Illumina NextSeq 2000 at the University of Michigan Advanced Genomics Core.

### CUTGRUN

E16 retinas from CD-1 mice were isolated, pooled, and dissociated into single cells using Accutase (Gibco). Approximately 3×10⁵ cells were used for each CUTCRUN assay, which were performed using the CUTCRUN Assay Kit (Cell Signaling Technologies, CST 86652S) following the manufacturer’s instructions for live cell preparation with modifications: micrococcal nuclease digestion was performed for 1hr in an ice-water bath (0° C), except for the Foxn4 samples and mouse IgG controls, which were digested with micrococcal nuclease overnight (∼12 hrs) at 0° C. Genomic DNA recovered from the CUTCRUN assay was used to prepare a DNA library using the NEBNext Ultra II DNA Library Prep Kit (NEB) for Illumina, adhering to the manufacturer’s protocol with the following modifications: after the addition of End Prep Buffer and Enzyme Mix, samples were incubated at 20°C for 30 minutes and subsequently at 50°C for 30 minutes. For ligation, 2.5 µl of 1:50 diluted NEB Unique Dual Index UMI Adaptors were added. Adaptor-ligated DNA was purified using 1.1X volume of NEBNext Sample Purification Beads. Following PCR enrichment, the adaptor-ligated DNA was again purified using a 1.1X volume of beads. The PCR product was separated on a 6% TBE gel (Invitrogen), and a gel slice corresponding to 200–500 bp (determined using size markers in an adjacent lane) was excised. The DNA from the gel slice was eluted, precipitated using 2-propanol (ThermoFisher), resuspended in Tris-HCl buffer (Invitrogen), and sequenced using an Illumina NextSeq 2000 at the University of Michigan Advanced Genomics Core.

### Bioinformatic analysis

Paired-end RNA-seq reads (∼55nt each) were mapped to mm39 using star (Dobin et al., 2013). Differential expression of genes in RNA-seq data was determined with DEseq2 (Love et al., 2014). For CUTCRUN analysis, ∼55nt paired-end Illumina sequences had adaptors removed using Cutadapt (Martin, 2011) and were then mapped to the mm10 mouse genome and to the Saccharomyces cerevisiae genome (for yeast spike-in DNA) with bowtie2 (Langmead and Salzberg, 2012). Mapped reads were deduplicated based on UMIs using UMI-Tools (Smith et al., 2017). Mapped paired-end reads were filtered for insert sizes of ≤120nt as recommended for transcription factor peak finding with CUTCRUN (Meers et al., 2019b; Meers et al., 2019c). Bedgraph files of read coverage for the mouse genome were generated using bedtools genomecov (Quinlan, 2014; Quinlan and Hall, 2010) and normalized based on total yeast spike-in read counts for each library. Peaks of read accumulation were identified using SEACR with relaxed peak finding (Meers et al., 2019a). Peaks were identified from two independent CUTCRUN libraries for each transcription factor, relative to two matched IgG control libraries. For Foxn3, peaks identified in two separate CUTCRUN experiments (each with two independent samples and two control samples) were intersected to generate a high confidence set of Foxn3 peaks that was used for subsequent Foxn3 binding region analysis.

The summit of each SEACR peak was defined as the center base in the region with the maximum read count. Peaks were filtered using exclude lists for CUTCRUN (Nordin et al., 2023) and ENCODE Chip-seq (Amemiya et al., 2019); non-chromosomal and chrM mappings also were discarded. Homer 5.1 (Heinz et al., 2010) was used to map DNA binding motifs in each peak. Peaks were linked to a gene annotation based on the nearest TSS from epdNewPromoter_v6.TSS (Dreos et al., 2012) for each peak summit, determined with bedtools closest. ENCODE cCREs (Abascal et al., 2020) that overlapped at least 10% of a SEACR peak were identified using bedtools intersect. Gene symbols for annotated CUTCRUN peaks were updated to current MGI symbols using https://www.informatics.jax.org/batch and matched with gene symbols in the RNA-seq data. Overlaps between peaks for different transcription factors were determined using bedtools intersect with at least 10% overlap.

Heat maps were generated using Deep Tools (Ramirez et al., 2016) and centered on peaks. Reads mapping at peaks were displayed with IGV (Thorvaldsdottir et al., 2013). Volcano plot was generated with VolcaNoseR (Goedhart and Luijsterburg, 2020). GO analysis was performed using NIH DAVID: https://david.ncifcrf.gov/tools.jsp (Huang da et al., 2009; Sherman et al., 2022). Venn diagrams were generated with the Broad Institute Venn diagram generator (http://barc.wi.mit.edu/tools/). Figures were assembled and labeled with Adobe Illustrator.

### Structure Predictions for Protein Interactions

Structure predictions for complexes of full-length proteins with DNA, or protein domains without DNA, were generated using AlphaFold 3: https://alphafoldserver.com (Abramson et al., 2024). See Supplemental Figure S9 for the sequences used. Structure predictions were visualized and interactions between proteins analyzed with The PyMOL Molecular Graphics System, Version 3.1.6.1 Schrödinger, LLC.

### Reporter Assays

HEK293 cells were cultured in 96-well plates in DMEM supplemented with 10% fetal bovine serum (FBS), 100 units/ml penicillin, and 100 µg/ml streptomycin (all from ThermoFisher). For each well, 20 ng of reporter DNA (Foxn3Enh2_WT, Foxn3Enh2_RfxM, Foxn3Enh2_FBM, Foxn3Enh2_EBM, Foxn3Enh1, Tek1, Ccdc39, Cfap206, Foxj1, Meis1, Notch1 or Ptprf) was co-transfected with 20 ng of either individual or combinations of the following DNA constructs: pUS2, pUS2-Foxn3, pUS2-Foxn3_7A, pUS2-Foxn4, pUS2-Foxn4_7A, pUS2-Foxj, pUS2-Ascl1, pUS2-Rfx3, pUS2-Rfx3-S616D, pUS2-Rfx3-F620A, pUS2-Rfx3-H624A, or pUS2-Rfx3-C544S. Additional pUS2 was added as needed to maintain the same total DNA amount. An additional 20 ng of pRL-SV40 was included as an internal control for transfection efficiency. Transfections were carried out using Lipofectamine 2000 (Invitrogen) or TransIT-VirusGEN (Mirus) per manufacturer’s instructions. Reporter activity was measured 24 hrs post-transfection using the Dual-Glo® Luciferase Assay System (Promega). Firefly luciferase activity was normalized to Renilla luciferase activity to account for transfection efficiency. Luminescence intensity was measured with a FLUOstar OPTIMA plate reader. The relative expression data shown in the figures include individual data points and the means for 3-6 independent transfections. Statistical analyses were performed using one-way or two-way ANOVA in GraphPad Prism 10.

### Co-immunoprecipitation and Western Blot Analyses

HEK293 cells in 10-cm dishes were co-transfected with 2 µg of each plasmid and 2 µg of the combination plasmid DNA using Lipofectamine 2000 or TransIT-VirusGEN (per manufacturer’s instructions). After 24 hrs, cells were lysed in NETN buffer (20 mM Tris-HCl, pH 8.0 (Invitrogen), 100 mM NaCl (Lonza), 0.5% NP-40 (Sigma), 0.5 mM EDTA (Invitrogen) supplemented with protease and phosphatase inhibitors (Sigma). Lysates were cleared by centrifugation (13,000 rpm, 5 min, 4 °C). For immunoprecipitation, 20 µL of Pierce™ Anti-HA Magnetic Beads (ThermoFisher) were incubated with lysates for 2 hrs at 4 °C with rotation, washed three times with NETN buffer, and eluted in 1× NuPAGE™ LDS sample buffer (Invitrogen) by boiling for 10 min. Proteins were resolved on 10% NuPAGE™ gels, transferred to nitrocellulose membranes (Invitrogen), and probed with rabbit anti-HA, mouse anti-Myc, or rabbit anti-Rfx3 (see Supplemental Table 5). Secondary antibodies included IRDye® 800CW goat anti-rabbit (1:8000, Li-COR), goat anti-rabbit StarBright™ Blue 700, and goat anti-mouse StarBright™ Blue 520 (1:5000, Bio-Rad). Blots were imaged using a Bio-Rad ChemiDoc™ MP system.

### Electrophoretic Mobility Shift Assay (EMSA)

30-bp oligonucleotides containing consensus Rfx3 and Foxn3 binding sites based on Foxn3 enhancer 2 were synthesized (IDT; see Supplemental Table 5), with or without a 5′-biotin-label, and annealed to create DNA duplex probes for EMSA. For Rfx3 binding, HEK293 cells in 6-cm dishes were transfected with 4 µg pUS2-MT-Rfx3 or point mutants using TransIT-VirusGEN. Protein extracts were prepared 24 hrs later by lysing cells in 30 µL of NETN buffer containing protease and phosphatase inhibitors. Lysates were cleared by centrifugation (13,000 rpm, 5 min, 4 °C). Rfx3 EMSA binding reactions (10 µL) contained 15 mM HEPES-KOH (pH 7.9, Gibco), 40 mM KCl (Sigma), 1 mM DTT (Invitrogen), 4% glycerol (Sigma), 50 ng/µL poly(dI-dC)(ThermoFisher), 400 fmol probe, and 2 µL extract, and were incubated at 4 °C for 30 min. For Foxn3 binding, HEK293 cells in 10-cm dish were transfected with 10 µg pUS2-HA-Foxn3 or pUS2-HA-Foxn3-7A. Extracts were prepared in NETN buffer, clarified, and HA-tagged Foxn3 proteins were purified using Pierce™ Anti-HA Magnetic Beads. The Foxn3 proteins were eluted with 20 µl of 2 mg/mL HA peptide (ThermoFisher) at 37 °C for 10 min. Foxn3 EMSA binding reactions (10 µL) containing 15 mM HEPES-KOH (pH 7.9), 40 mM KCl, 1 mM DTT, 5 mM MgCl₂ (Ambion), 400 fmol probe, and 3 µL purified Foxn3, were incubated at room temperature for 30 min. For competition assays, 40 pmol of unlabeled competitor oligonucleotide duplex was added before the labeled probe. Complexes were resolved on a 6% native polyacrylamide gel (ThermoFisher) in 0.5× TBE (Invitrogen) at 100 V for 70 min at 4 °C, transferred to a nylon membrane (Amersham) overnight at 50 V and 4 °C, UV-crosslinked, and blocked for 15 min with 1× blocking buffer (250 mM NaCl, 17 mM NaH₂PO₄, 7.3 mM NaH₂PO₄, 1% SDS (Sigma)). Membranes were incubated with streptavidin–HRP (Vector Laboratories) in blocking buffer for 10 min, washed three times with 0.1× blocking buffer (5 min each), and visualized by chemiluminescence using Clarity Max™ Western ECL substrates (Bio-Rad).

## Acknowledgements

We are grateful to Jillian Pearring for helpful discussions and comments on the manuscript, antibodies, and assistance with microscopy. We thank Amy Liu for assistance with cell counting. We thank Ray Trievel for advice and discussions about AlphaFold and predicted protein structures. We thank Anne Vojtek and Mike Uhler for helpful discussions and comments on the manuscript. We thank Andrew Tidball and Jack Parent for assistance with microscopy and use of the Andor BC43.

## Data Availability

Sequencing data are available through NCBI GEO: accessions GSE306960 and GSE306964.

