## Supplemental Figures for "*Foxn3* is part of a transcriptional network that regulates primary cilia in the developing retina"

*Supplemental Figures S1-S16*

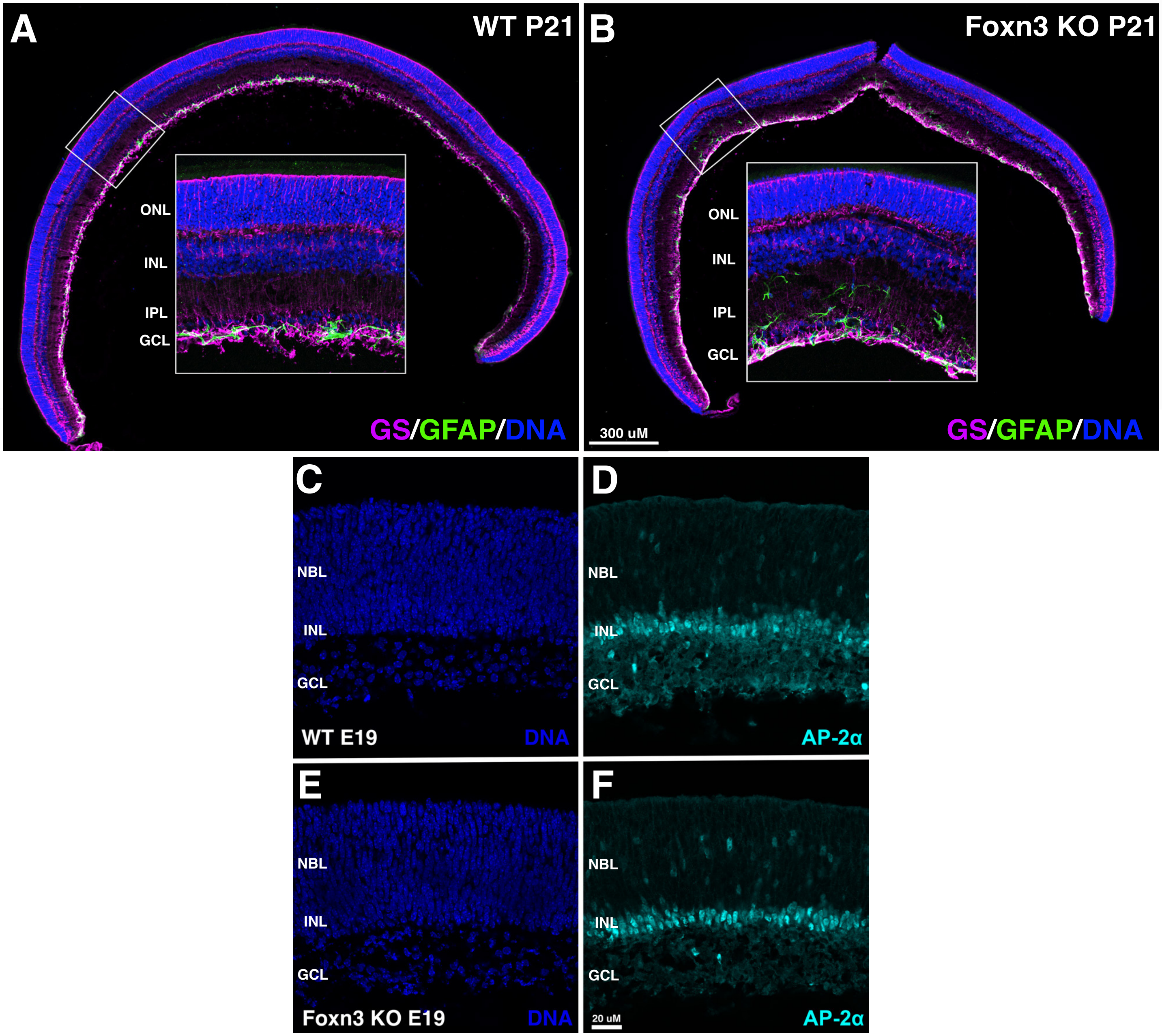


**Figure S1**. Glial cells labeled by GFAP (green) or glutamine synthase (GS, magenta) in P21 retina sections. (A, B) GS-positive Müller glial cell bodies are present in the INL in both WT and *Foxn3* KO retinas. GFAP-positive astrocytes are restricted to the GCL and inner IPL of WT retinas, while GFAP-positive astrocytes are present in the GCL and throughout the IPL of *Foxn3* KO retinas. At E19, sections from WT (C, D) or *Foxn3* KO (E, F) retinas do not show obvious differences either in overall structure or in the location of amacrine cells detected by AP2α immunofluorescence (cyan). DNA/nuclei: blue.

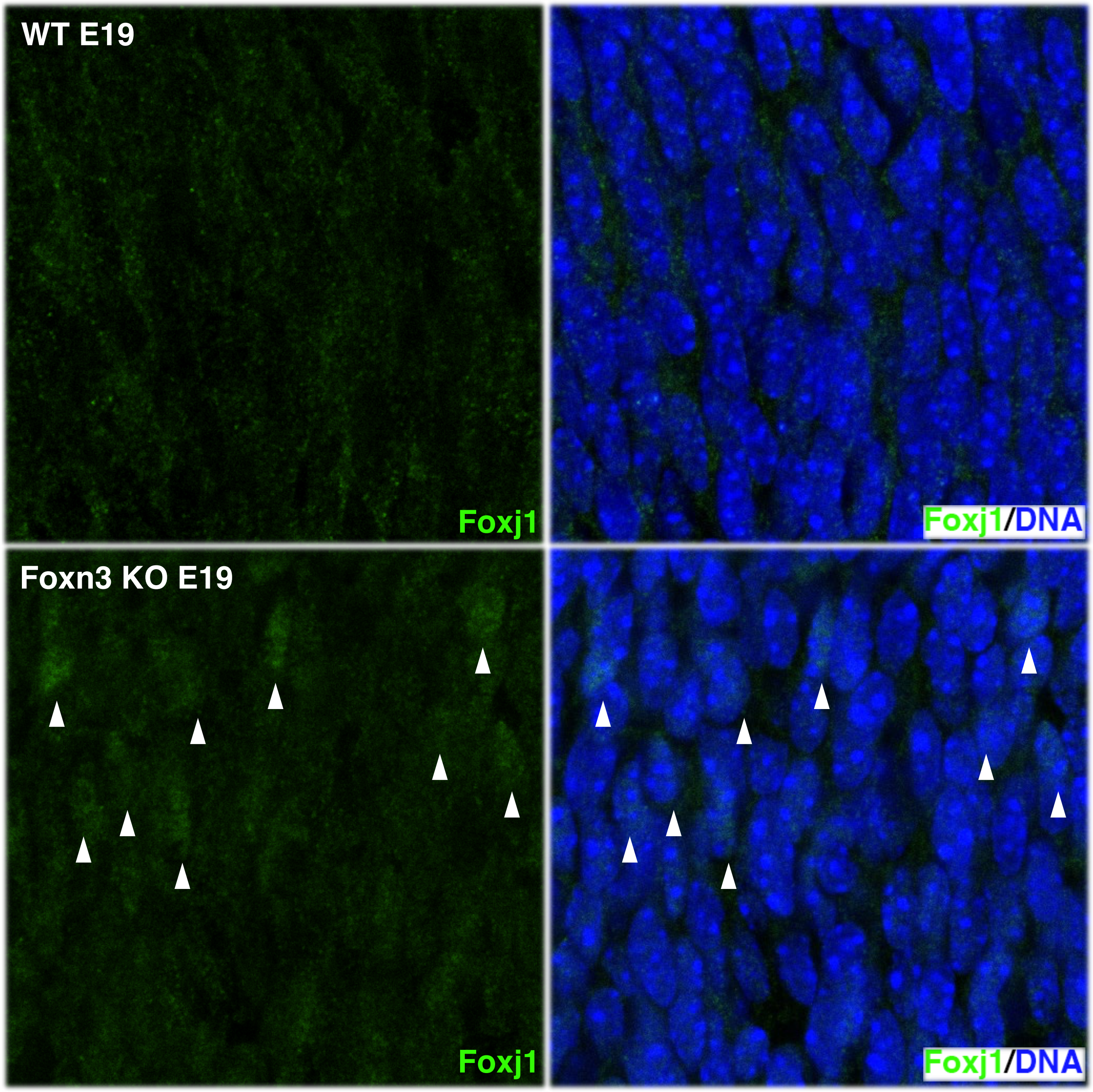


**Figure S2.** Little or no Foxj1 protein (green) is detected in the neuroblast layer of WT retinas at E19 by immunofluorescence (A, B). Increased Foxj1 protein is detected in a subset of nuclei (arrowheads) in the neuroblast layer of E19 retinas from the *Foxn3* KO (C, D).

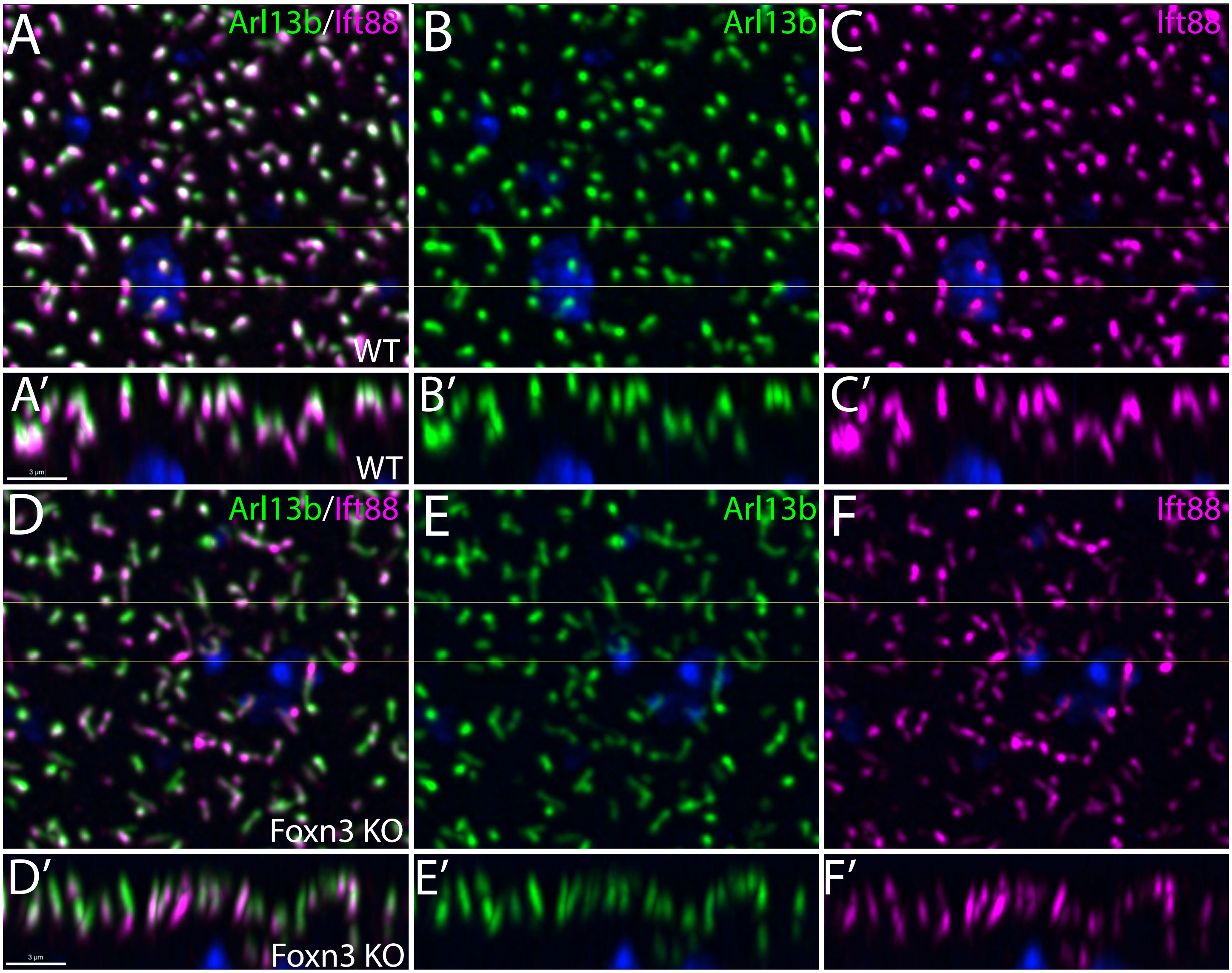


**Figure S3.** Altered primary cilia in the developing retina in the absence of *Foxn3*. (A-C) Immunofluorescence detection of Arl13b (green) and Ift88 (magenta) in primary cilia at the outer surface of a representative whole-mount E19 WT retina. Maximum intensity projection of top 7μm of the retina (Z axis). DNA/nuclei: blue. (A'-C') Maximum intensity projection for 3μm virtual transverse section (Y axis) through the retina cilia; the boundaries of the section are indicated by yellow lines in A-C. (D-F) Primary cilia at the outer surface of a representative whole-mount E19 *Fox3* KO retina, with a virtual transverse section (D'-F'), details as for A-C/A'-C'. Primary cilia are disorganized and have more variable localization of Ift88 within cilia for the *Foxn3* KO retina compared to the WT retina. Scale bars (3μm) in A' and D' apply to all panels.

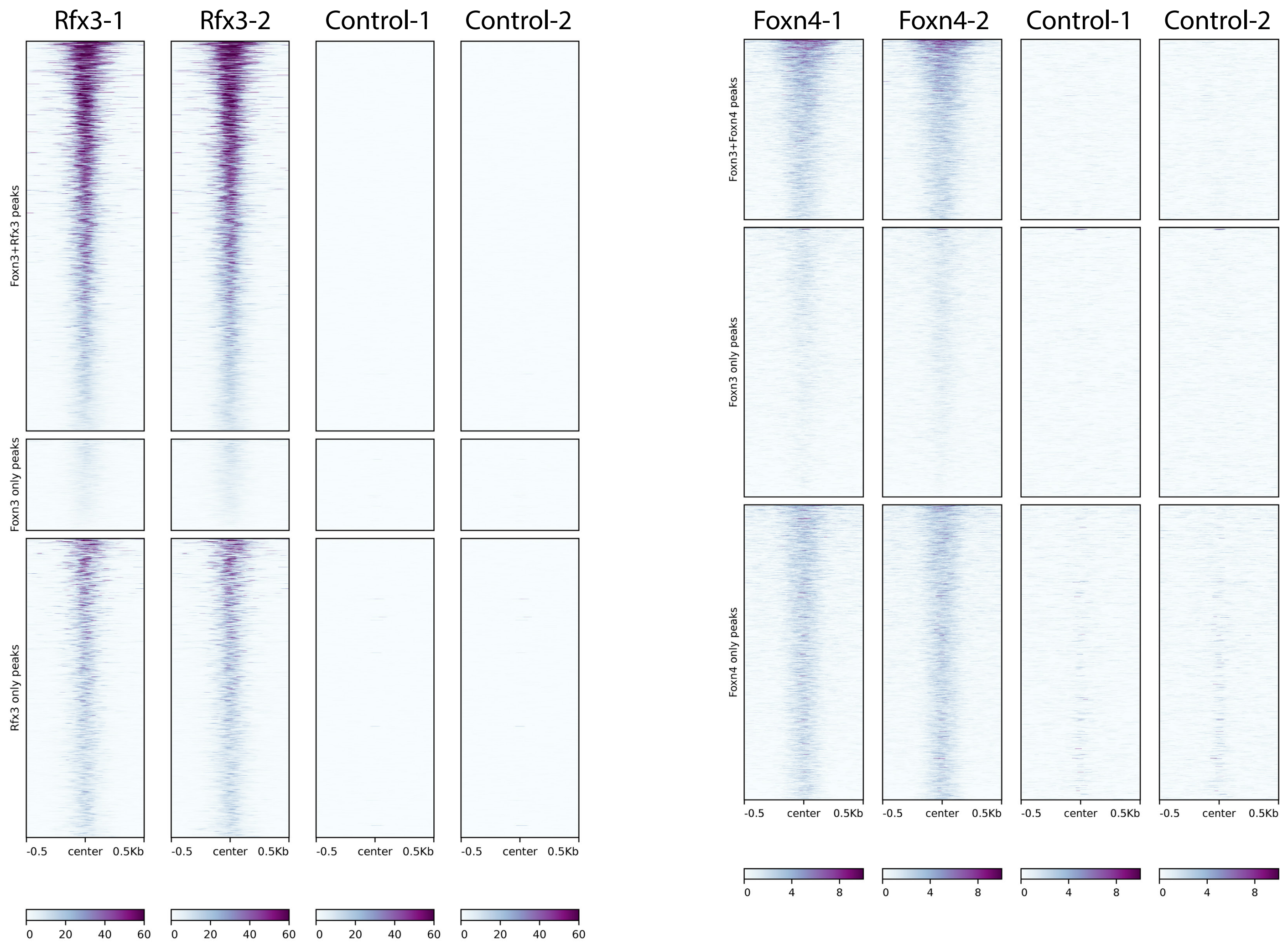


**Figure S4.** Heat maps showing reproducibility of individual CUT&RUN samples for Rfx3 and Foxn4. Left: Rfx3 and a matching control antibody. Right: Foxn4 and a matching control antibody. Labels are the same as for Figure 5.

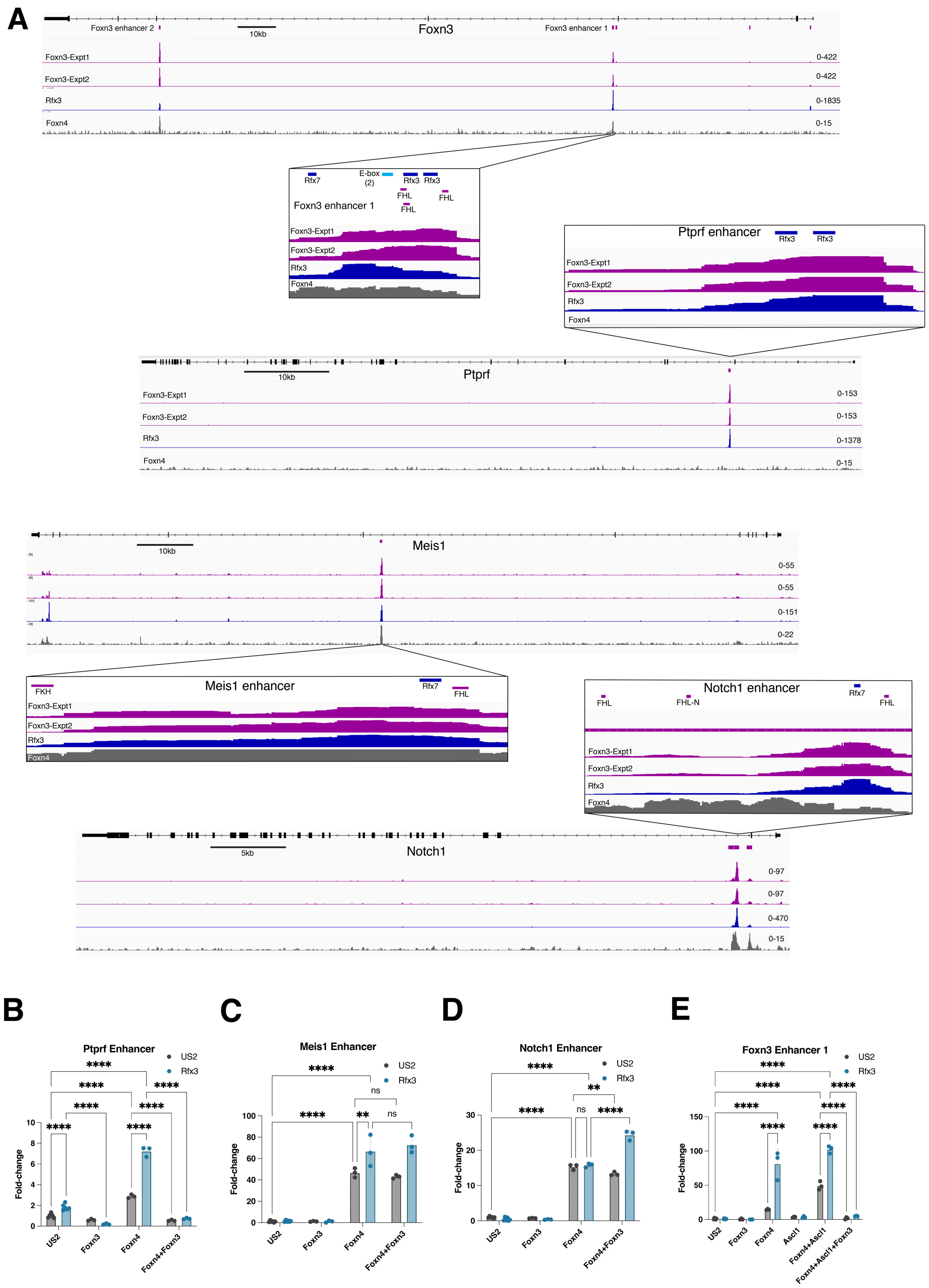
A
Foxn3 enhancer 2
10kb
Foxn3
Foxn3 enhancer 1
Foxn3-Expt1
0-422
Foxn3-Expt2
0-422
Rfx3
0-1835
Foxn4
0-15
Rfx7
E-box
(2)
Rfx3
Rfx3
FHL
FHL
FHL
Foxn3 enhancer 1
Foxn3-Expt1
Foxn3-Expt2
Rfx3
Foxn4
Ptprf enhancer
Rfx3
Rfx3
Foxn3-Expt1
Foxn3-Expt2
Rfx3
Foxn4
10kb
Ptprf
Foxn3-Expt1
0-153
Foxn3-Expt2
0-153
Rfx3
0-1378
Foxn4
0-15
10kb
Meis1
0-55
0-55
0-151
0-22
Meis1 enhancer
FKH
Rfx7
FHL
Foxn3-Expt1
Foxn3-Expt2
Rfx3
Foxn4
Notch1 enhancer
FHL
FHL-N
Rfx7
FHL
Foxn3-Expt1
Foxn3-Expt2
Rfx3
Foxn4
5kb
Notch1
0-97
0-97
0-470
0-15
B
Ptprf Enhancer
US2
Rfx3
Fold-change
US2
Foxn3
Foxn4
Foxn4+Foxn3
C
Meis1 Enhancer
US2
Rfx3
ns
ns
Fold-change
US2
Foxn3
Foxn4
Foxn4+Foxn3
D
Notch1 Enhancer
US2
Rfx3
ns
Fold-change
US2
Foxn3
Foxn4
Foxn4+Foxn3
E
Foxn3 Enhancer 1
US2
Rfx3
Fold-change
US2
Foxn3
Foxn4
Ascl1
Foxn4+Ascl1
Foxn4+Ascl1+Foxn3

**Figure S5.** Regulation of reporters based on regulatory elements bound by Foxn3, Rfx3, and/or Foxn4 in four different genes. (A) The *Foxn3*, *Ptprf1*, *Meis1*, and *Notch1* genes contain intronic enhancers bound by both Foxn3 and Rfx3 based on CUT&RUN data from E16 embryonic retinas. Foxn4 binding in the E16 retinas was detected by CUT&RUN at the enhancers for *Foxn3*, *Meis1*, and *Notch1*, but not *Ptprf1*. The location of consensus binding sites for forkhead, RFX, and bHLH factors are indicated in the enlarged regions. The *Ptprf* enhancer does not contain a consensus forkhead binding site, although it does contain two Rfx3 binding sites within the region bound by both Foxn3 and Rfx3. (B-E) Luciferase reporters based on the four enhancers were constructed and assayed by transfection into HEK293 cells in combination with Foxn4, Foxn3, and/or Rfx3 expression vectors. Foxn4 activated all reporters and cooperatively activated with Rfx3 on all reporters except *Meis1*, although activation of the *Ptprf* reporter was modest. The *Foxn3* enhancer 1 reporter (E) also was tested with a co-transfected Ascl1 expression vector, which further increased activation in combination with Foxn4 or Foxn4 and Rfx3. Co-expression of Foxn3 repressed activation of the *Ptprf1* reporter and the *Foxn3* enhancer 1 reporter regardless of the activating proteins. In contrast, Foxn3 did not prevent activation of the *Meis1* or *Notch1* reporters. N=3-6 for reporter assays, 2-way ANOVA: **** $p \leq 0.0001$, ** $p \leq 0.01$, ns = not significant.

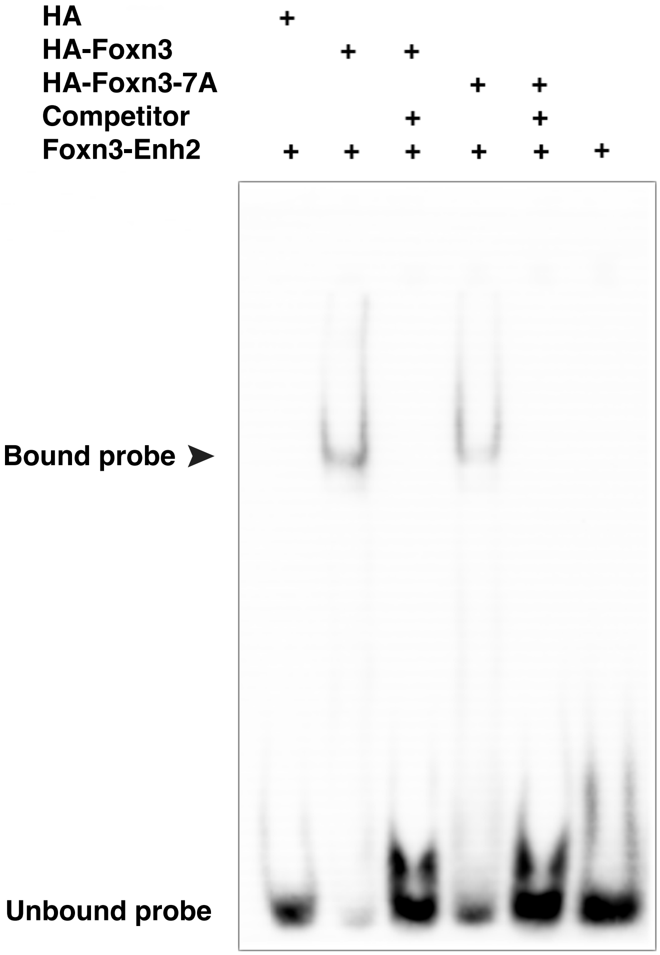


**Figure S6.** Both HA-Foxn3 and HA-Foxn3-7A proteins bind to DNA. The HA-Foxn3 and HA-Foxn3-7A proteins were purified from transfected HEK293 cells and tested for in vitro DNA binding with EMSA, using a biotinylated 30nt double-stranded DNA probe containing FKH and Rfx3 sites from the Foxn3 enhancer 2 (Foxn3-Enh2. See Fig.6A and Supplemental Table 5). Both proteins shifted the DNA probe (arrowhead), and binding was competed by unlabeled DNA probe (Competitor). HA indicates purified protein from a control vector expressing only the HA-epitope tag.

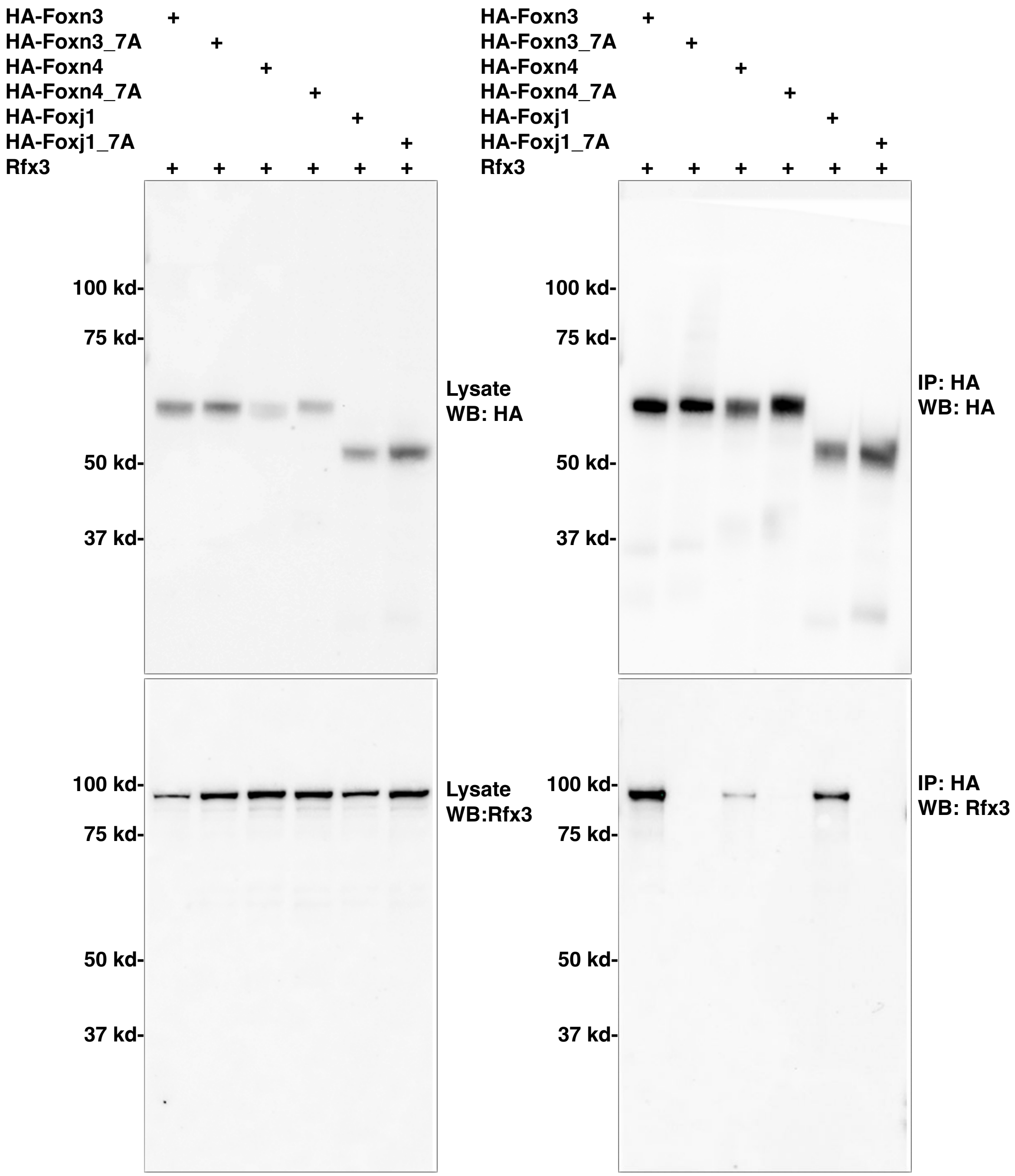


**Figure S7.** Complete images for western blots shown in Fig. 7B.

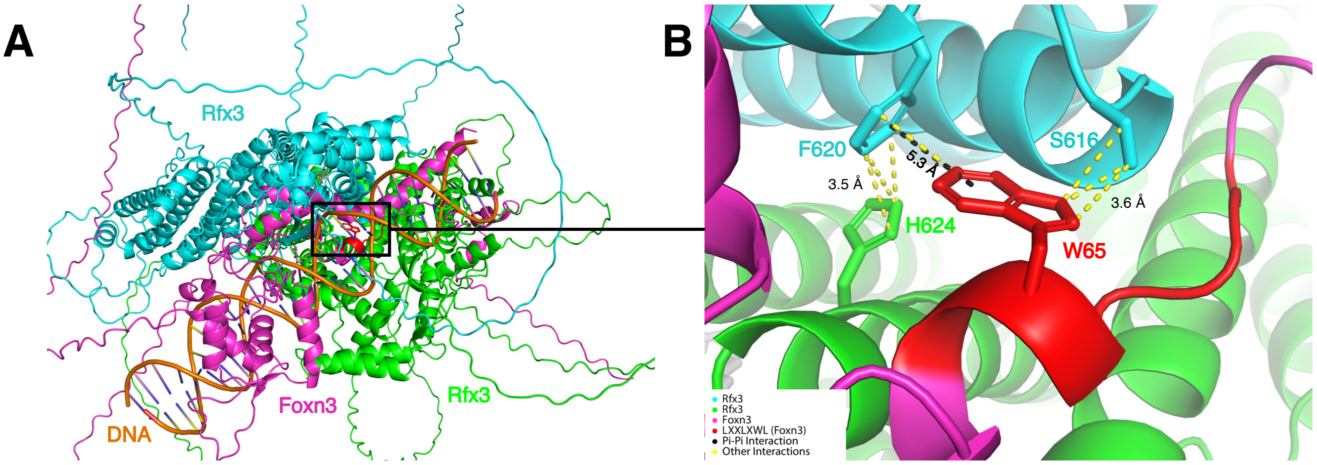


C

| *Model* | *pTM* | *ipTM* |
|---|---|---|
| Full Length Rfx3 - Full Length FoxN3 and DNA | 0.51 | 0.46 |
| Rfx3 Dimer - FoxN3 N-terminus | 0.86 | 0.82 |
| Rfx3 Dimer - FoxN4 N-terminus | 0.84 | 0.79 |
| Rfx3 Dimer - FoxJ1 N-terminus | 0.84 | 0.80 |

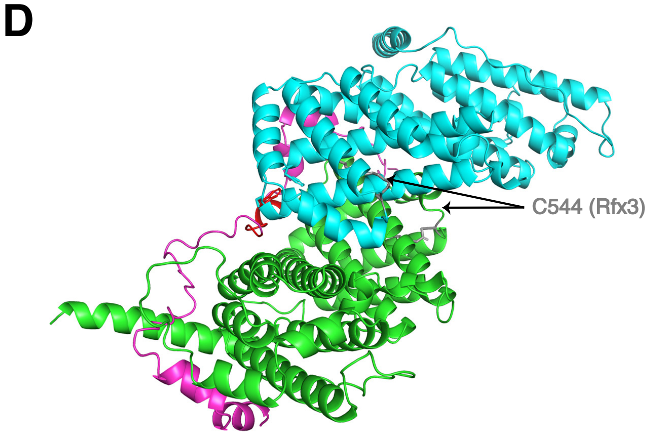


**Figure S8.** AlphaFold 3 predicted interaction between an Rfx3 dimer and a Foxn3 protein bound to DNA. A dimer of Rfx3 (Full Length Protein) interacting with Foxn3 (Full Length Protein) and a DNA molecule bound by both proteins was modeled with AlphaFold 3 (see Figure S7 for sequences). Both Rfx3 molecules are palmitoylated at their C544 residues (gray). Other colors according to legend. (A) Full view of the Rfx3 (two molecules), FoxN3, and DNA predicted interaction with each protein/DNA labelled. (B) Zoomed-in image of the interaction site. Tryptophan (W65) from FoxN3 is labelled along with serine (S616), phenylalanine (F620), and histidine (H624) residues from Rfx3. T-shaped pi-pi interaction (black) between F620 of Rfx3 and W65 of FoxN3 with a distance of 5.3 Å. Other interactions (yellow) between all four amino acids are indicated, of varying distances. (C) Table of AlphaFold3 prediction confidence scores. ipTM values above 0.8 are considered high-quality predictions. (D) Alternate view of the Rfx3 B, C, and D domain dimer with the FoxN3 N-terminus showing the palmitoylated C544 residue (gray) on the opposite side of the dimerization domain from the predicted interaction with Foxn3.

| ***Protein/DNA*** | ***Sequence used for AlphaFold 3 models*** |
|---|---|
| Rfx3 C-terminus<br>(B, C, and D Domains) | GSGQQTGTSVEQTVIAQSQHHQQFLDASRALPEFGEVEISSLPDGTTFEDIKSLQSLYRE<br>HCEAILDVVVNLQFSLIEKLWQTFWRYSPSTPADGTTITESSNLSEIESRLPKAKLITLCKH<br>ESILKWMCNCDHGMYQALVEILIPDVLRPIPSALTQAIRNFAKSLEGWLSNAMNNIPQRMI<br>QTKVAAVSAFAQTLRRYTSLNHLAQAARAVLQNTSQINQMLSDLNRVDFANVQEQASW<br>VCQCDDNMVQRLETDFKMTLQQQSTLEQWAAWLDNVMMQALKPYEGRPSFPKAARQ<br>FLLKWSFYSSMVIRDLTLRSAASFGSFHLIRLLYDEYMFYLVEHRVAQVTGETPIAVMGEF<br>GDLN<br>= Palmitoylated C544 |
| FoxN3 N-Terminus | GLSQRYRGSGFSKALQEDDDLDFPLPDIRLEEGAMEDEELTNLNWLHESKNLLKSFGES<br>VLRSVSPVQDLDDDTPPS |
| FoxN4 N-Terminus | MIESGIWSRMSEMIRSSGHSHHCSPQEYRFLPPVGDDDLPGDLQSLSWLTAVDVPRLQ<br>QMANGRIDLGSSGVTHPHP |
| FoxJ1 N-Terminus | MAESWLRLCGAGPGEEAGPEGGMEEPDALDDSLTSLQWLQEFSILNAKAPTLPPGGTD<br>PHGYHQVPGLVAPGSPLAADP |
| Rfx3 Full Length Protein | MQTSETGSDTGSTVTLQTSVASQAAVPTQVVQQVPVQQQVQQVQTVQQVQHVYPAQV<br>QYVEGSDTVYTNGAIRTTTYPYTETQMYSQNTGGNYFDTQGSSAQVTTVVSSHSMVGT<br>GGIQMGVTGGQLISSSGGTYLIGNSMENSGHSVTHTTRASPATIEMAIETLQKSDGLSTH<br>RSSLLNSHLQWLLDNYETAEGVSLPRSTLYNHYLRHCQEHKLDPVNAASFGKLIRSIFMG<br>LRTRRLGTRGNSKYHYYGIRVKPDSPLNRLQEDMQYMAMRQQPMQQKQRYKPMQKV<br>DGVADGFTGSGQQTGTSVEQTVIAQSQHHQQFLDASRALPEFGEVEISSLPDGTTFEDI<br>KSLQSLYREHCEAILDVVVNLQFSLIEKLWQTFWRYSPSTPADGTTITESSNLSEIESRLP<br>KAKLITLCKHESILKWMCNCDHGMYQALVEILIPDVLRPIPSALTQAIRNFAKSLEGWLSN<br>AMNNIPQRMIQTKVAAVSAFAQTLRRYTSLNHLAQAARAVLQNTSQINQMLSDLNRVDF<br>ANVQEQASWVCQCDDNMVQRLETDFKMTLQQQSTLEQWAAWLDNVMMQALKPYEGR<br>PSFPKAARQFLLKWSFYSSMVIRDLTLRSAASFGSFHLIRLLYDEYMFYLVEHRVAQVTG<br>ETPIAVMGEFGDLNAVSPGNLDKDEGSEVESETDEDLDDSSEPRAKREKTELSQAFPVG<br>CMQPVLESAVQPSLLNPLHSEHIVTSTQTIRQCSATGNTYTAV<br>= Palmitoylated C544 |
| FoxN3 Full Length Protein | MGPVMPASKKAESSGISVSSGLSQRYRGSGFSKALQEDDDLDFPLPDIRLEEGAMEDEE<br>LTNLNWLHESKNLLKSFGESVLRSVSPVQDLDDDTPPSPAHSDMPYDARQNPNCKPPY<br>SFSCLIFMAIEDSPTKRLPVKDIYNWILEHFPYFANAPTGWKNSVRHNLSLNKCFKKVDKE<br>RSQSIGKGSLWCIDPEYRQNLIQALKKTPYHPPPTPQAYQSTSGPPIWPGSTFFKRNGAL<br>LQVSPGVIQNGARVLSRGLFPGVRPLPITPIGMTAAIRNSITSCRMRTESEPPCGSPVVS<br>GDPKEDHNYSSAKSSTARSTSPTSDSISSSSSSADDHYEFATKGSQEGSEGSFQSHES<br>HSEPEEEDRKPSPKEGKDALGDSGYASQHKKRQHFAKARKVPSDTLPLKKRRTEKPPE<br>SDDEEMKEAAGSLLHLAGIRSCLNNITNRTAKGQKEQKETAKN |
| DNA sequences for Rfx3-FKH binding site from FoxN3 enhancer 2 (double stranded) | 5’ GAAGGTCACCATGGCAACGGTAAACAACCC 3’<br><br>5’ GGGTTGTTTACCGTTGCCATGGTGACCTTC 3’ |

**Figure S9**. Mouse protein and DNA sequences used for AlphaFold 3 structure predictions shown in Fig. 8A, B and in Supplemental Figures S6 and S8. The Rfx3 sequence was palmitoylated at C544 for the AlphaFold 3 predictions.

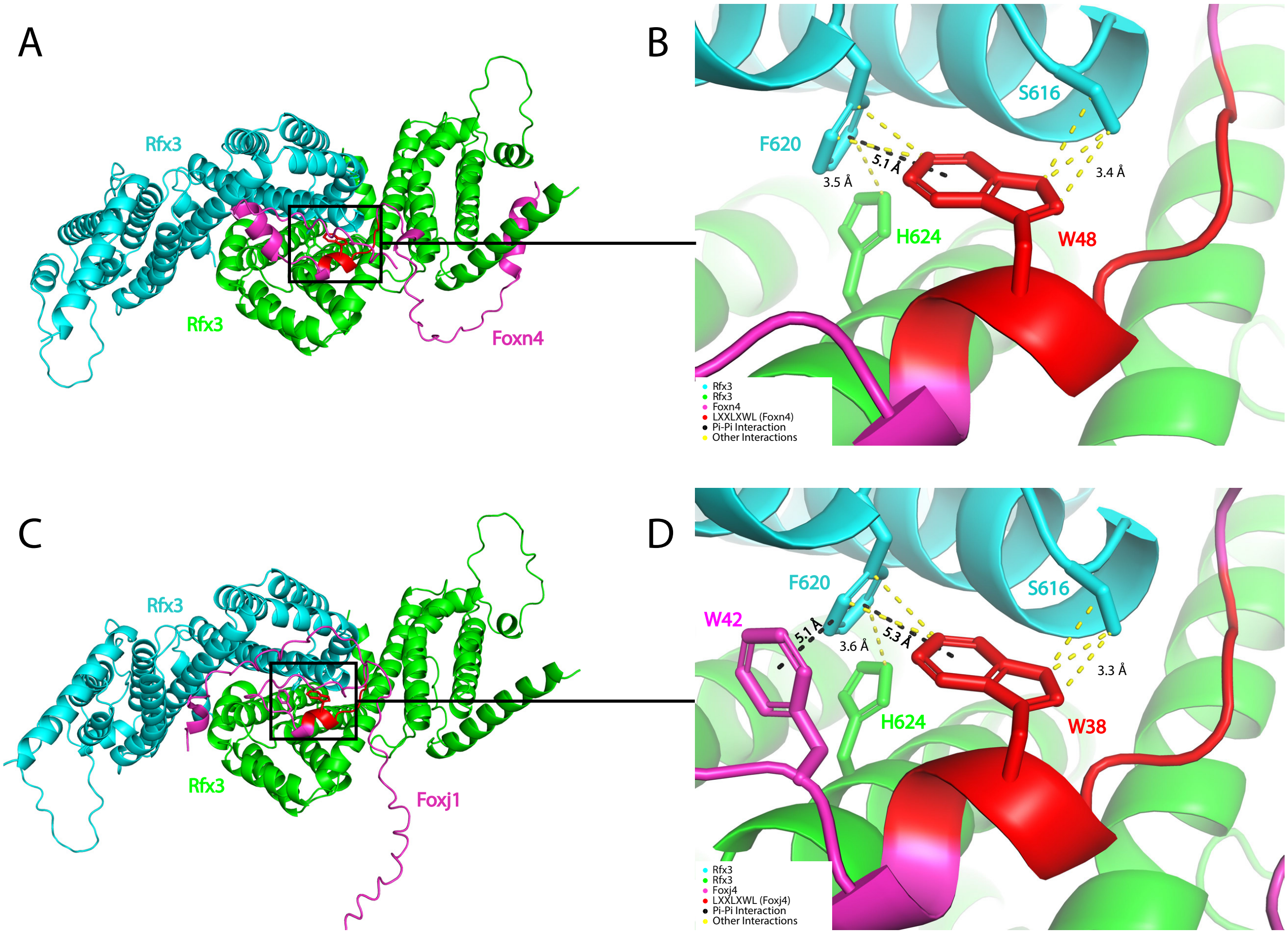


**Figure S10.** AlphaFold 3 predicted interactions between Rfx3 and Foxn4 or Foxj1 protein domains are similar to the predicted interaction between Rfx3 and Foxn3 protein domains (Fig. 8A, B). Interactions between a homodimer of the Rfx3 B, C, and D domains and the Foxn4 N-terminus or Foxj1 N-terminus were modeled by AlphaFold 3. Rfx3 molecules are palmitoylated at C544 residues. Colors according to legend. (A) Full view of Rfx3 dimerization domain (two molecules) and Foxn4 N-terminus interaction with each protein labelled. (B) A zoomed-in image of the interaction site. Tryptophan W48 from Foxn4 labeled along with serine, phenylalanine, and histidine residues from Rfx3. A pi-pi interaction is shown (black) between F620 of Rfx3 and W48 of Foxn4 with a distance of 5.1 Å. (C) Full view of Rfx3 dimerization domain (two molecules) and Foxj1 N-terminus interaction with each protein labeled. (D) A zoomed-in image of the interaction site. Tryptophan W38 from Foxj1 labeled along with serine, phenylalanine, and histidine residues from Rfx3. Two pi-pi interactions are shown (black), between F620 of Rfx3 and W38 of Foxj1 with a distance of 5.3 Å, and between F620 of Rfx3 and W42 of Foxj1 with a distance of 5.1 Å.

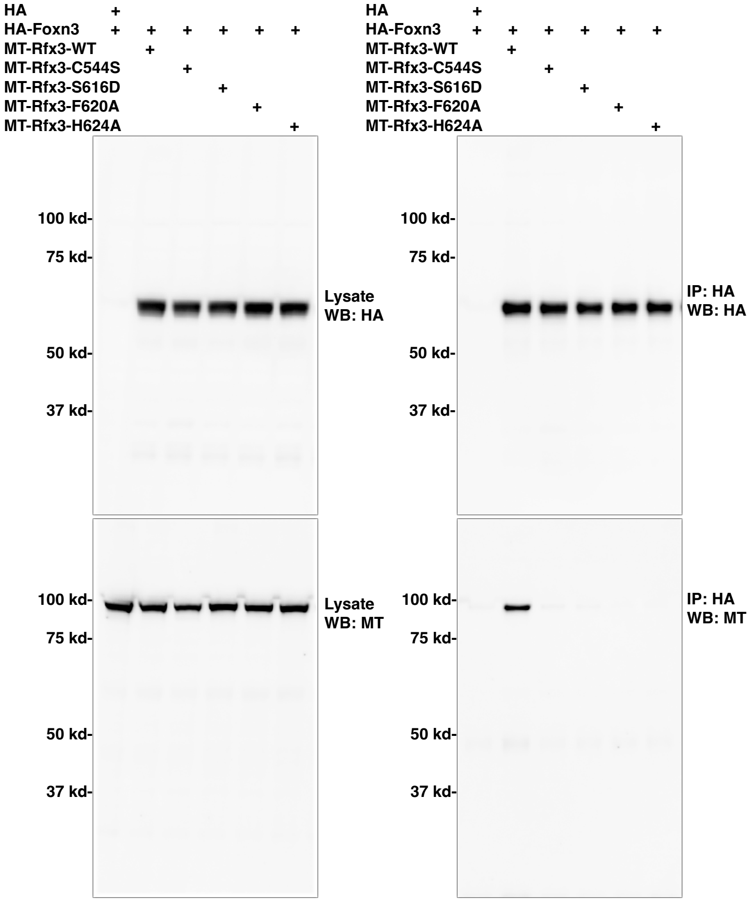


**Figure S11.** Complete images for western blots shown in Fig. 8C.

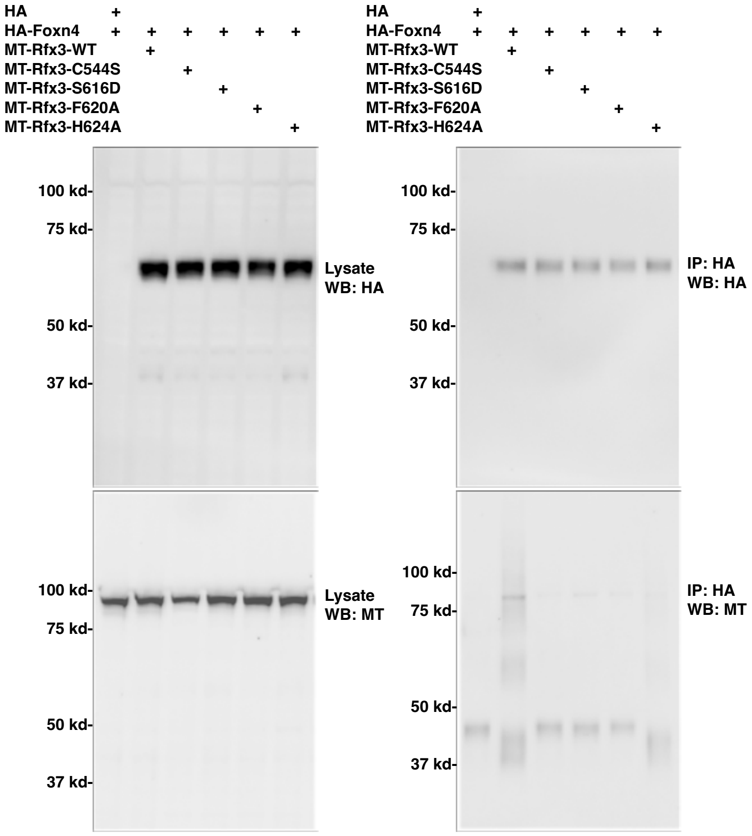


**Figure S12.** Complete images for western blots shown in Fig. 8D.

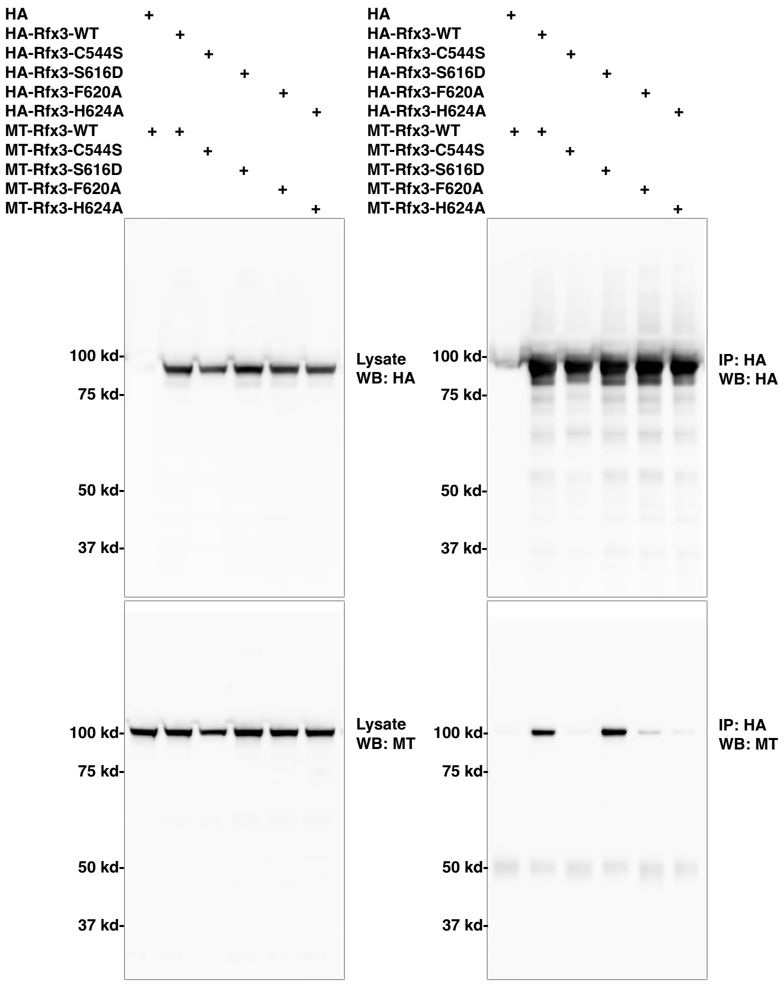


**Figure S13.** Complete images for western blot data shown in Fig. 8F.

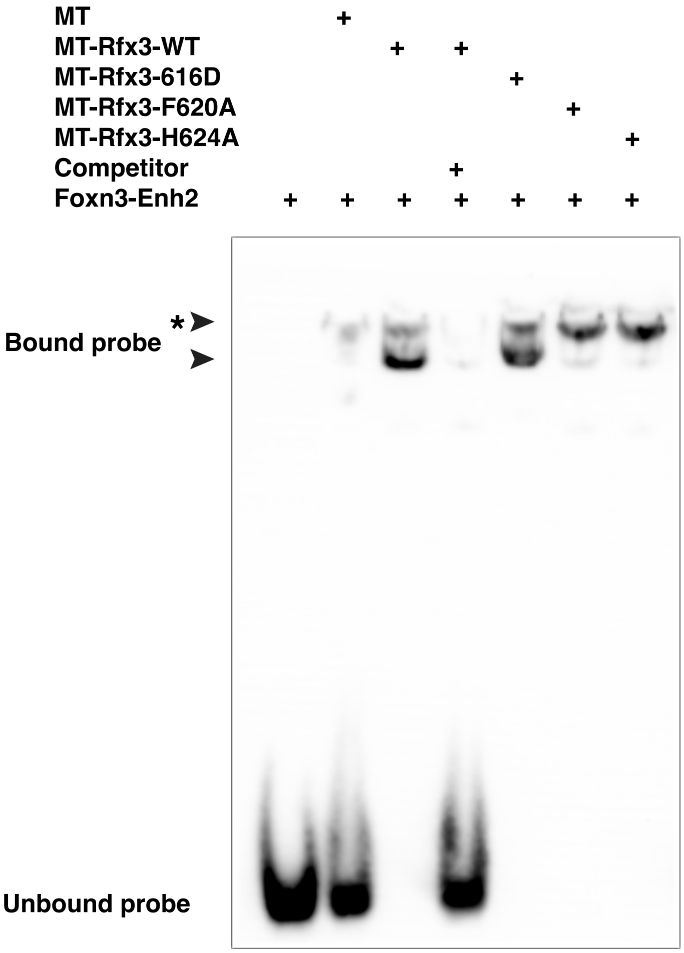


**Figure S14.** Rfx3 mutant proteins bind DNA. Expression vectors for Rfx3 with 6 Myc-epitope tags at the N-terminus (MT-Rfx3) or each of three different MT-Rfx3 point mutant proteins were transfected into HEK293 cells and protein extracts were tested for in vitro DNA binding with EMSA, using a biotinylated 30nt double-stranded DNA probe containing FKH and Rfx3 sites from the Foxn3 enhancer 2 (Foxn3-Enh2, see Fig.6A and Supplemental Table 5). All proteins bound to DNA and shifted either one or two major bands (arrowheads). MT-Rfx3 and MT-Rfx3 S616D generated two shifted bands, while the MT-Rfx3 F620, and MT-Rfx3 H624 proteins generated only the upper shifted band (*). The lower shifted band may be generated by binding of an Rfx3 dimer (see text). First lane shows probe without HEK293 extract. MT indicates probe with HEK293 protein extract for a control vector expressing only the Myc-tag. MT-Rfx3 binding was competed by unlabeled probe DNA (Competitor).

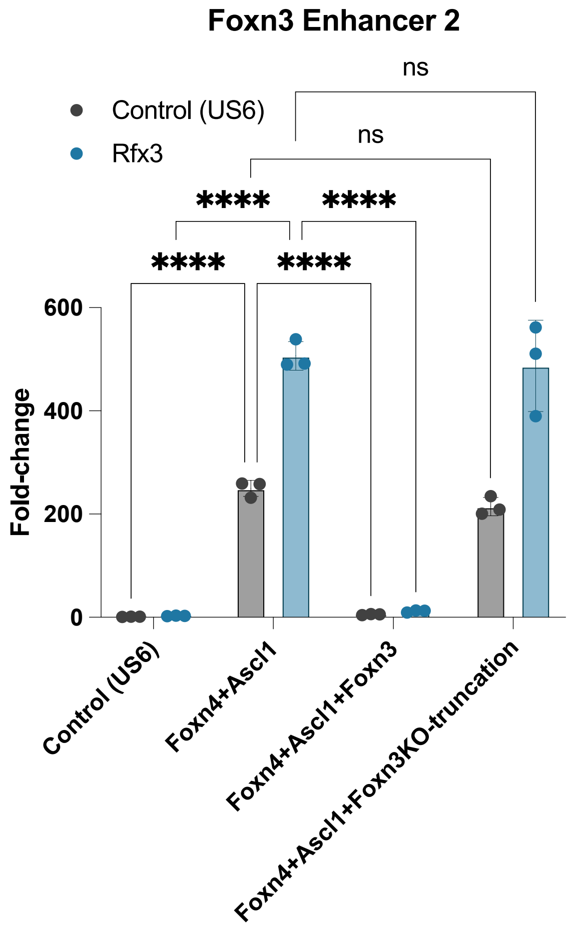


**Figure S15.** A truncated Foxn3 protein that is potentially expressed in *Foxn3* KO mice is not functional in a reporter assay. The Foxn3 gene in the Foxn3 KO mouse has a deletion of exon 3. Exon 2, containing the Foxn3 initiation codon and N-terminal coding region (including the LXXLXWL motif and part of the forkhead domain), can be spliced to exon 4, leading to a truncated Foxn3 protein followed by a novel tail from out-of-frame translation of part of the C-terminal coding region. The Foxn3KO-truncation vector expresses the Foxn3 coding region without exon 3. Co-expression of Foxn3 represses activation of the reporter by Foxn4 with Ascl1, with or without co-expression of Rfx3. However, co-expression of the Foxn3KO-truncation has no effect on activation by Foxn4 with Ascl1, with or without co-expression of Rfx3, suggesting that the potential truncated Foxn3 protein in the *Foxn3* KO is non-functional. N=3. 2-way ANOVA: **** p≤0.0001, ns = not significant.

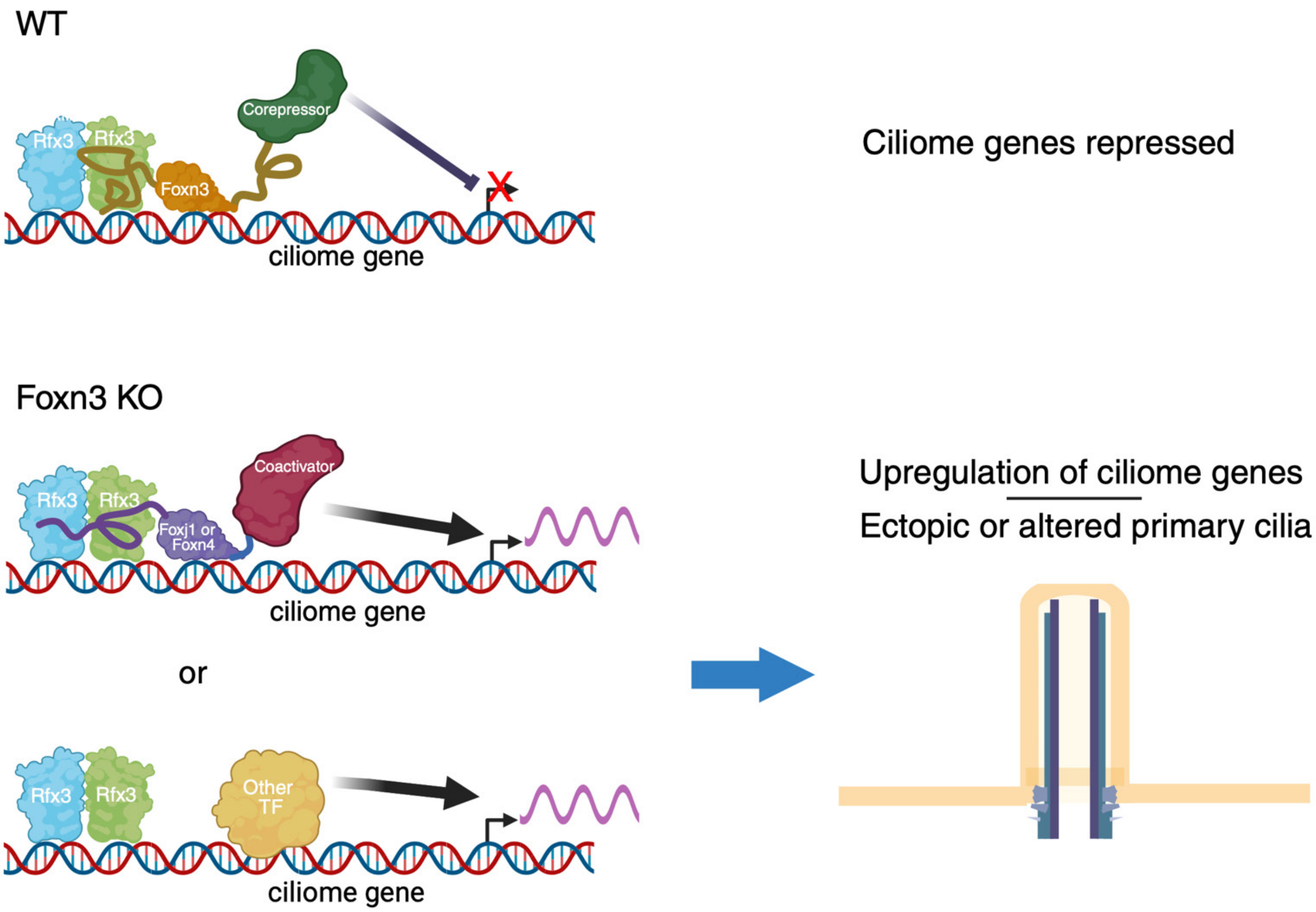


**Figure S16.** Model of Foxn3 function in the retina. In the developing WT retina, Foxn3 prevents or limits expression of genes in the ciliome in a subset of cells. Foxn3 binds at specific target sites and recruits corepressors to repress transcription of target genes. Most Foxn3 genomic binding sites in the embryonic retina are also bound by the Rfx3 protein. The Foxn3 LXXLXWL motif interacts with the Rfx3 dimerization domain in an Rfx3 protein dimer and that interaction is required for Foxn3 repression. Foxn4 and Foxj1 also contain the LXXLXWL motif and interact with Rfx3 to cooperatively activate transcription of target genes. Since Foxn3 can bind to the same target sequences as Foxj1 or Foxn4, it can compete with these proteins for DNA binding and for interaction with Rfx3, in addition to recruiting corepressors, likely enhancing target gene repression. Foxn3, Foxn4, or Foxj1 also can interact with Rfx3 dimers in solution (not shown). In *Foxn3* KO retinas, loss of the Foxn3 protein allows increased expression of Foxj1 and other ciliome genes, activated either by Foxj1/Foxn4 or by other transcription factors. Increased ciliome gene expression allows formation of ectopic cilia on retinal bipolar or amacrine interneurons, as well as possible alterations of primary cilia on other cells.

Fig. S16 created in BioRender. Turner, D. (2025) https://BioRender.com/8u8r2o4.
